# A synthetic ERFVII-dependent circuit in yeast sheds light on the regulation of early hypoxic responses of plants

**DOI:** 10.1101/2025.09.01.673502

**Authors:** Mikel Lavilla-Puerta, Yuming He, Luca Piccinini, Lorenzo Di Paco, Antonis Papachristodoulou, Francesco Licausi, Beatrice Giuntoli

## Abstract

Plants face hypoxic conditions either chronically, as particular tissues are characterized by fluctuating or stable low oxygen levels, or acutely, when flooded. In vascular plants, transcriptional adaptive responses to hypoxia are rapidly mounted by Ethylene Response Factors VII (ERFVIIs), regulated by Plant Cysteine Oxidases (PCOs) through the cysteine branch of the N-degron pathway (Cys-NDP) for oxygen sensing. However, this relatively simple regulatory circuit, consisting of both constitutively expressed as well as hypoxia-inducible ERFVIIs and PCOs, interacts with diverse signalling cues and pathways invoked by hypoxia. To understand the share of the PCO-mediated oxygen sensing mechanism in the production of hypoxia responses, we insulated the PCO/ERFVII circuit from *Arabidopsis thaliana* and adapted it to *Saccharomyces cerevisiae*. Using a reporter gene to monitor the output of the circuit allowed us to compare the speed and amplitude of response to hypoxia in the engineered yeast and the source organism. Hypoxia triggered ERFVII stabilization both in Arabidopsis and yeast, leading to a similarly fast transcriptional response that was however larger in plants. A simple hypoxia-inducible feedback loop improved the amplitude of response in yeast, demonstrating the importance of this regulation in the endogenous PCO/ERFVII circuit. Finally, computational modelling of the yeast circuit enabled us to identify promoter competition and presence of hypoxia-inducible PCOs as key parameters that shape early hypoxia responses in plant cells.

**Significance Statement:** We report the design, testing and optimisation of a synthetic molecular switch that activates gene expression in response to hypoxia in the yeast *Saccharomyces cerevisiae*. This is based on enzymes that consume molecular oxygen to regulate the stability of transcription factors in plant cells. By generating such a hybrid molecular device, we were able to demonstrate the efficacy of this hypoxia response strategy independently of the many ancillary components that affect gene regulation in plant cells. In this way, we were able to assess its activation dynamics, characterised by similarly fast induction of gene expression in both yeast and plants. Our approach also revealed the requirement of interlocked feedback loops to achieve the magnitude of gene induction measured in plants.

## Introduction

The enrichment of Earth’s atmosphere with molecular oxygen dramatically changed the planet’s biogeochemistry and likely impacted on the diversification of life forms (1). Aerobic organisms use this compound for several essential metabolic reactions, including cellular respiration, although they are regularly exposed to fluctuations in oxygen levels (2). Therefore, all aerobes have developed mechanisms to monitor the availability of oxygen and initiate transcriptional responses when its levels fall below their metabolic requirements (hypoxia). Fungi indirectly rely on the abundance of metabolites whose synthesis requires O_2_, like sterols or heme (3, 4). Plants and animals converged instead on transcription factors (TFs) whose abundance is post-translationally controlled in an O_2_-dependent fashion (2).

In flowering plants, the response to hypoxia is largely mediated by TFs belonging to the group VII of Ethylene Response Factors (ERFVIIs) (5, 6). ERFVII families from Angiosperms present a variable number of members that, while all participating to low oxygen responses, differ in terms of hypoxia-inducibility and appear to be partially not redundant in activity (5, 7, 8). Most ERFVII proteins are subjected to a common mechanism for O_2_-dependent regulation, controlled by the proteolytic N-degron pathway (9, 10). Their N-terminal cysteine, exposed upon co-translational methionine cleavage by Met aminopeptidases (MetAP) (11) is aerobically oxidized into its sulfinyl (CysO_2_) or sulfonyl form (CyO_3_) by Plant Cysteine Oxidases (PCOs) (12, 13). Oxidation in turn stimulates N-terminal arginine conjugation by arginyltransferase enzymes (ATEs) and subsequent polyubiquitination of proximal Lys by ubiquitin ligases (PRT6 and BIG) (14), before the ERFVIIs are degraded by the proteasome. This series of reactions constitutes the so-called PCO- or Cys-branch of the N-degron pathway, which also enables oxygen perception in metazoan animals (15).

Beyond ERFVII conditional O_2_-dependent degradation, additional mechanisms associated with hypoxia impact on the stability or activity of these TFs. First, mitochondrial retrograde signalling promotes ERFVII stabilisation and the consequent activation of hypoxia responsive genes (16). Second, fatty acid desaturation favours ERFVII detachment from the plasma membrane and their relocation to the nucleus (17, 18), in cells where they are protected from degradation. Reduced ATP levels under hypoxia have been connected to a shift of fatty acid composition towards oleoyl-CoA in Arabidopsis seedlings (19). Finally, it has been found that ERFVII phosphorylation in response to early Ca^2+^ signalling or TOR-mediated energy signalling stimulate stabilisation and transactivation capacity (20, 21).

Recently, a subset of ER-tethered NAC transcription factors has been connected to low-oxygen stress responses in *Arabidopsis thaliana* (22, 23). Among them, ANAC013 relocates to the nucleus in the initial phases of hypoxia and associates to core anaerobic gene promoters (24) by contacting their cognate DNA element. Although their role is yet to be fully understood, ANACs have been proposed to cooperate with the ERFVII in the regulation of hypoxia-inducible genes, in response to mitochondrial disfunction signals (25).

The network of signalling mechanisms invoked by hypoxia, together with the genetic redundancy of their components, raises the question as to whether the cellular response to hypoxia is directly initiated by a decrease in oxygen levels, or, alternatively, it requires the cooperation between the O_2_-sensing pathway and indirect signalling of hypoxia. It also remains to be defined how much the transcriptional adjustments to hypoxia can be explained in terms of PCO-mediated ERFVII activity and how much additional pathways contribute to this process. To address these fundamental questions, we resorted to the principles of synthetic biology (26). We reconstructed the ERFVII-dependent transcriptional regulation *de novo* in baker’s yeast (*Saccharomyces cerevisiae*), a chassis that promises to minimize the crosstalk between the synthetic circuit and endogenous regulation. Indeed, yeast oxygen perception operates via heme- and ergosterol-dependent pathways (3, 4) and no PCO or ERFVII homologues exist (27). We have previously demonstrated that a plant-inspired Cys N-degron pathway can be established in yeast (27); here, we introduced the PCO/ERFVII signalling module to regulate gene expression.

Abstraction of plant modules, in a cellular context unhindered by concurrent regulatory mechanisms, provided us with the unprecedented opportunity to measure the contribution of the Cys N-degron pathway to the initiation and maintenance of the hypoxic response. Moreover, we could use the quantitative output data from the synthetic yeast to build a model summarizing the properties of a minimal PCO/ERFVII oxygen-dependent circuit. This approach complements the information on the features of plant oxygen sensing obtained so far, through the *in vitro* characterization of PCO enzymes (12) and *in planta* assessment of gene dynamics (5).

## Results

### Plant cells rapidly mount fast transcriptional responses to acute hypoxia

The onset of low oxygen conditions is known to determine broad reconfigurations in plant transcriptomes (28, 29), however the information about the promptness of these responses is scant. To tackle the earliest events of hypoxic signalling, we exposed fully aerated *Arabidopsis thaliana* seedlings to acute low oxygen (1% O_2_ atmosphere) and monitored the expression of hypoxia markers in the first minutes of stress (5’, 10’, 15’), as well as after more prolonged treatment (1 h, 2 h). Transcripts of all nine hypoxia-responsive genes (HRGs) we selected (24) increased significantly above aerobic levels within 5 to 10 minutes and kept accumulating over the treatment, following logistic curves (**Fig. 1A, SI Appendix, Table S1**). To quantify the speed of activation of each HRG, we used the response time (RT), or the time required to increase mRNA levels from 10% to 90%. *HRA1* (*Hypoxic Response Attenuator 1*) and *LBD41* (*Lateral organ Binding Domain 41*) transcripts increased most rapidly. Two other genes also involved in hypoxia signalling, *PCO1* (*Plant Cysteine Oxidase 1*) and *HRE2* (*Hypoxia Responsive ERF 2*), had intermediate speed of induction, while the rest of the markers, including the hypoxic metabolism landmarks *PDC1* (*Pyruvate Decarboxylase 1*), *ADH1* (*Alcohol Dehydrogenase 1*) and *PGB1* (*Phytoglobin 1*), increased more progressively (**Fig. 1A**).

**Figure 1.**
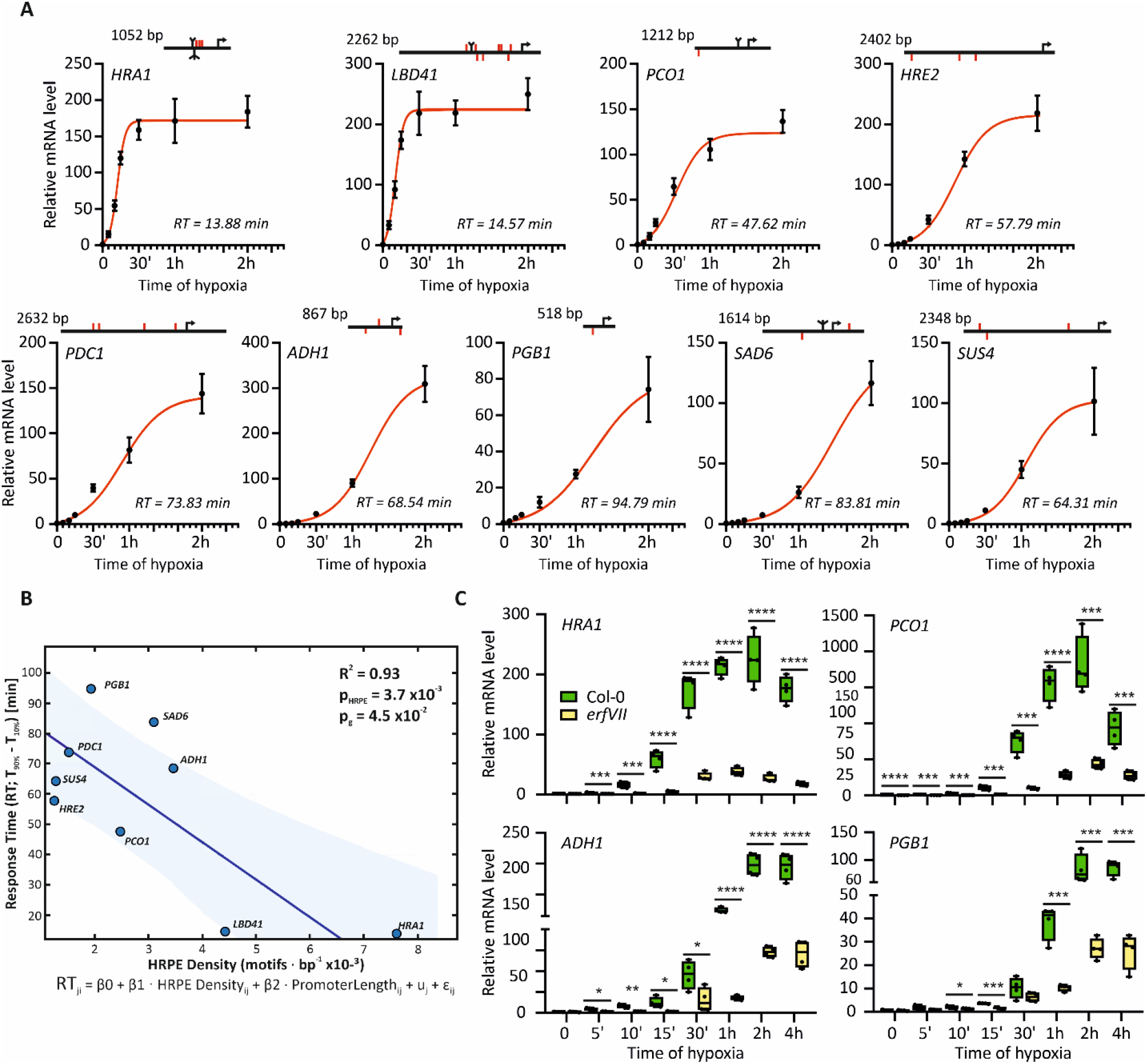
Short-term profiling of HRG expression in *A. thaliana* seedlings. **(A)** Time-resolved expression of four hypoxia responsive genes: *Hypoxic Response Attenuator 1* (*HRA1*), *Lateral organ Binding Domain 41* (*LBD41*), *Plant Cysteine Oxidase 1* (*PCO1*), *Hypoxia Responsive ERF 2* (*HRE2*), *Pyruvate Decarboxylase 1* (*PDC1*), *Alcohol Dehydrogenase 1* (*ADH1*), *Phytoglobin 1* (*PGB1*), *Stearoyl-Acyl Carrier Protein 6* (*SAD6*) and *Sucrose Synthase 4* (*SUS4*) in 7-day-old Col-0 seedlings growing on vertical plates moved from normoxia (t_0_) to hypoxia (1% O_2_ v/v) for 2 h. RT, response time. Above, architecture of the same genetic loci, spanning from up to 2 kb upstream of the ATG codon (arrow) down to the first intron of the gene. Individual HRPE elements are highlighted in red, while bi- or tridents indicate 2 or 3 overlapping motifs, respectively. HRPE orientation is also displayed with marks above (presence in the sense DNA strand) or below the horizontal line (presence in antisense strand). **(B)** Correlation between response time (RT) and HRPE density (motifs bp^-1^) in the HRGs from (A). Equation below (see SI Appendix, Extended methods for details on the linear model). **(C)** Expression of selected markers in Col-0 and *erfvii* mutant seedlings undergoing an extended 4 h hypoxic treatment, similar to (A). Statistically significant differences between Col0 and *erfVII* after Student’s t-test are indicated by asterisks (*, 0.01≤p<0.05, **, 0.001≤p<0.01, ***, 0.0001≤p<0.001, ****, p<0.0001; n=4).

To understand the contribution of the ERFVII factors to HRG fast activation, we looked at the distribution of hypoxia-responsive promoter elements (HRPE), the ERFVII cognate *cis-*motif (30), in their upstream regions (**Fig. 1A**). Using a linear mixed-effects model (SI Appendix), HRPE motif density in HRG promoters was strongly negatively associated with response time (*β*_*1*_ = −12.4 min per motif/kb; SE = 2.69 min per motif/kb; *p* = 0.0037), indicating faster responses at higher HRPE density (**Fig. 1B**). The scaled promoter length showed a modest negative association (*β*_2_ = −13.82 min; SE = 5.48 min; *p* = 0.045) while the intercept was *β*_*0*_ = 94.93 min (SE = 9.40 min; *p* = 5.48×10^-5^). Genes with faster response also showcased more concentrated HRPE sequences (**Fig. 1A**). In the pentuple *erfVII* mutant (31), the immediate induction of all markers was abolished (**Fig. 1C, SI Appendix, Fig. S1**), indicating that no other factor could replace the ERFVIIs as trigger of hypoxic gene activation. At later time points, a partial response was slowly mounted by the *erfVII*. This observation is compatible with the previous report of residual *ERFVII* expression (32).

We monitored ERFVII protein dynamics in early hypoxia, taking advantage of transgenic plants that express the ERFVII RELATED TO APETALA2 3 (RAP2.3) tagged at the C-terminus with a triple HA epitope (33). The extent of post-translational regulation we observed was comparable with existing reports (5, 33). Proteasome inhibition with bortezomib (BZ) led to strong RAP2.3 accumulation, while hypoxia had milder effect (**Fig. 2A, B** and **SI Appendix, Fig. S2A**). This could be due to a repression of translation (34), or to alternative mechanisms that concur to ERFVII degradation when the Cys-NDP is inhibited, as reported in the *prt6* mutant before (5). Closer inspection revealed that RAP2.3^3xHA^ stabilization occurred as early as 5 minutes into hypoxia and further protein accumulation was visible until 30 minutes (**Fig. 2C, D** and **SI Appendix, Fig. S2B**). RAP2.3^3xHA^ levels tended to decline slowly later, over 8 h of treatment. In contrast with RAP2.3, a direct target of O_2_-dependent proteolysis, the hypoxia-inducible proteins ADH and PDC showed a delayed accumulation in hypoxia (**SI Appendix, Fig. S2C**). These data overall support the conclusion that RAP2.3^3xHA^ stabilization by hypoxia occurs before the induction of HRGs in Arabidopsis seedlings.

**Figure 2.**
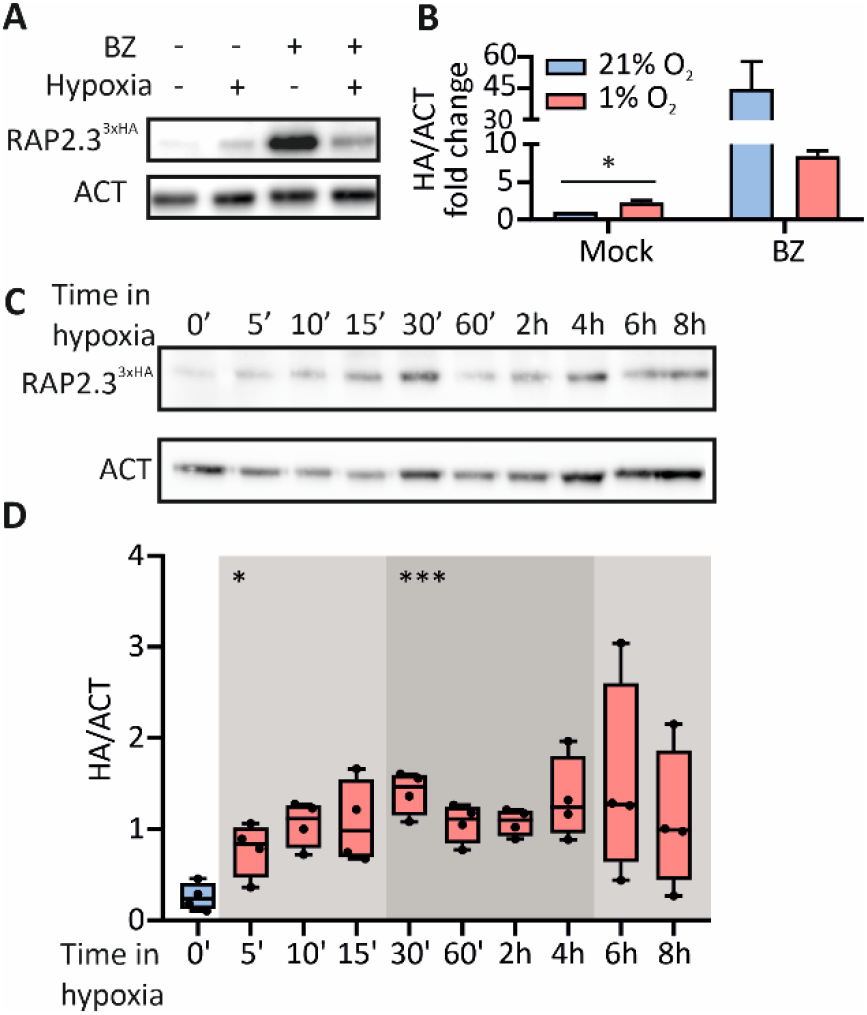
ERFVII protein regulation in short-term hypoxia. **(A)** Detection of RAP2.3^3xHA^ (HA) and actin (ACT) in liquid grown *35S:RAP2*.*3*^*3xHA*^ seedlings, after 6 h treatment with normoxia, 1% O_2_, 100 µM bortezomib (BZ), or an equal amount of DMSO (1% v/v, mock). **(B)** Densitometric quantification of RAP2.3^3xHA^ band intensity from the previous blots, normalized to actin (mean+SD, n=2). Data are expressed as fold change from the mean normoxic mock value. **(C)** RAP2.3^3xHA^ and actin abundance in 7-day-old seedlings from vertical plates, over a time-course of hypoxia extended up to 8 h. **(D)** Quantification of RAP2.3^3xHA^ band intensity (HA/ACT signal ratio), performed as in (B). Full blots are provided in **SI Appendix, Fig. S2**. Asterisks in (B) indicate statistically significant differences after Student’s t-test (*, 0.01≤p<0.05; n=4), shadings and asterisks in (D) represent statistically significant differences from normoxia from normoxia (T_0_), for One-way Anova (*, 0.01≤p<0.05, ***, 0.0001≤p<0.001; n=4).

### A plant-inspired switch induces gene expression under hypoxia in baker’s yeast

The previous profiling demonstrates that what enables seedlings to perceive and react to sudden hypoxia within minutes is ERFVII enrolment and stabilisation preceding activation of the transcriptional response. We hereby propose that the inhibition of the proteolytic Cys-NDP pathway can be the all-sufficient cause for the immediate onset of hypoxic transcription. To challenge our hypothesis, we decided to insulate the Cys-NDP from plant cells and test the dynamics of the hypoxic response into an orthogonal biological system (*S. cerevisiae*). Our plant-inspired circuit for hypoxia-controlled transcription consists of the Cys-NDP reduced to its essential components: a PCO sensor, an ERFVII effector and a *promoter:luciferase* reporter module. The combination of these three modules was expected to generate minimal crosstalk with endogenous regulatory pathways.

Nanoluciferase (NLUC) was chosen for the reporter module, by virtue of its intense signal and fast turnover (35). To confer full orthogonality to the circuit, we adopted a synthetic promoter made of tandem repetitions of the conserved 30-bp HRPE *cis-*element (30, 36). We tested three hypoxia-responsive HRPE promoter versions (HRPE, HRPE_ADH_ and HRPE_Ω_) that differ in their 5’ untranslated region (37). Once integrated in the TRP locus of the yeast strain W303, the promoters showed different levels of basal expression (**SI Appendix, Fig. S3A**). When compared with a normalizing construct, driven by the constitutive P*GPD* promoter from *glyceraldehyde 3-phosphate dehydrogenase*, HRPE- and HRPE_Ω_-NLUC produced 10- or 60-fold higher signal, respectively. In contrast, HRPE_ADH_-NLUC produced a low basal output, displaying a desirable feature for inducible systems.

We used the ERFVII protein RAP2.12 from Arabidopsis to design the effector module. The plant TF alone proved unable to activate any HRPE reporter (**Fig. 3A**). We thus tested RAP2.12 C-terminal fusion with three distinct transcription activation domains (TAD): the native yeast GAL4 AD and two synthetic TADs, 6TAL-VP64 and GAL4STE12 (**SI Appendix**, Extended methods). We hereby obtained the synthetic transcription factors RAP2.12-GAL4AD, RAP2.12-6TVP and SYRAP (RAP2.12-GAL4STE12, or Synthetic Yeast RAP). Moreover, fusion of SYRAP with an N-terminal ubiquitin monomer generated UbSYRAP (**SI Appendix**, Extended methods). This TF exposes an N-terminal Cys residue upon ubiquitin-mediated protein cleavage, a technique exploited before on a model Cys-NDP substrate in yeast (27).

**Figure 3.**
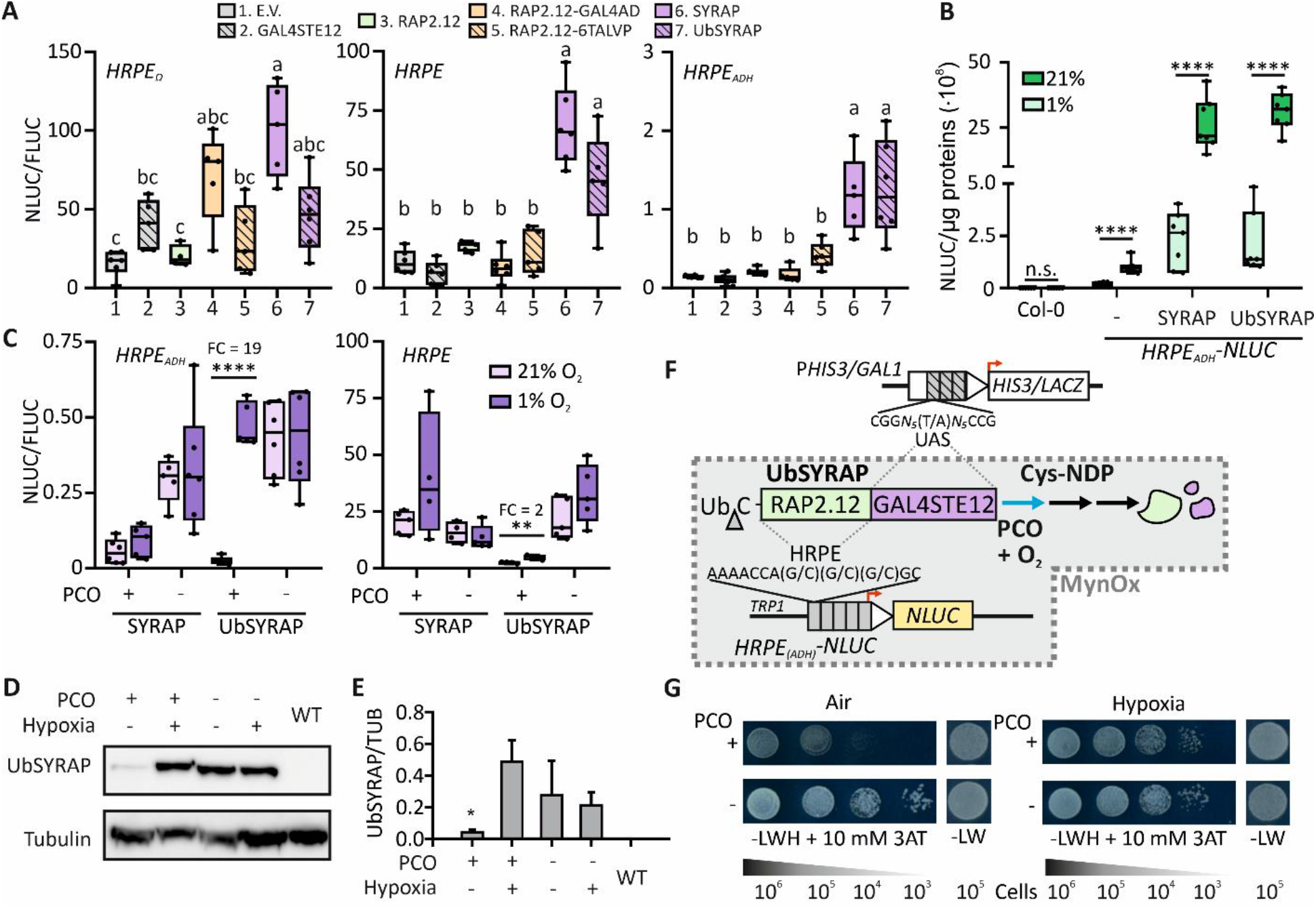
Assembly of orthogonal ERFVII-based transcriptional circuits in yeast. **(A)** Activation of three synthetic promoter versions (*HRPE-, HRPE*_*ADH*_*- and HRPE*_*Ω*_*-NLUC*) by RAP2.12-derived TFs in W303 cultures. Cells were co-transformed with a P*GPD-FLUC* integrative construct and HRPE promoter activity was expressed as relative NLUC activity (NLUC/FLUC) (n=5-6). E.V., empty vector. **(B)** Transient expression of *35S:SYRAP* constructs in agroinfiltrated leaves of Arabidopsis plants. The activity of an *HRPE*_*ADH*_*-NLUC* reporter, stably integrated in the genome, was recorded after 6 h incubation in normoxia or hypoxia (n=6). Water-infiltrated leaves from the wild type (“Col-0”) or *HRPE*_*ADH*_*-NLUC* (“-”) were included. **(C)** O_2_-dependent regulation of *HRPE-* and *HRPE*_*ADH*_*-NLUC* reporters in W303 yeast expressing SYRAP or UbSYRAP along with PCO or a control protein (GUS, −). Cultures (n=4-5) were incubated for 6 h in normoxia or hypoxia (1% O_2_). NLUC activity was normalized to P*GPD-FLUC* activity. Fold changes values (hypoxia vs. air; FC) are reported above the significantly different groups. **(D)** Regulation of UbSYRAP protein abundance, detected with an anti-GAL4BD antibody, by PCO and hypoxia (6 h, 1% O_2_). WT, aerobic untransformed cells. **(E)** Relative UbSYRAP amount to α-tubulin (mean + SD, n=3), quantified by densitometry from blots in **SI Appendix, Fig. S3C. (F)** Schematic overview of the synthetic O_2_-responsive circuits. The RAP2.12- and GAL4-derived DNA binding domains of UbSYRAP make the chimeric TF able to recognize both HRPE and UAS_GAL_ elements. UbSYRAP acts as effector of *NLUC* expression in W303 strains, or *HIS3* and *LACZ* expression in MaV203 strains. O_2_ enables UbSYRAP degradation through PCO, thereby lowering the output(s). The wedge indicates ubiquitin (Ub) cleavage by yeast deubiquitinases. **(G)** Growth of MaV203 cells expressing UbSYRAP in combination with PCO (+) or GUS (-), after 3 days incubation in normoxia or hypoxia. SD –LW, non-selective medium; SD –LWH, selective medium, supplemented with 10 mM 3-AT. The approximate number of cells spotted is reported below. Different letters in (A) or asterisks in (E) indicate statistically significant differences (p<0.05) after One-way Anova; asterisks in (B) and (C) indicate statistically significant differences after Student’s t-test on pairwise comparisons (*, 0.01≤p<0.05, **, 0.001≤p<0.01, ****, p<0.0001).

SYRAP and UbSYRAP outperformed the other two chimeric TFs. GAL4 AD and 6TAL-VP64 domains were not effective when fused with RAP2.12 (**Fig. 3A**). Failure of the first domain may have arisen from a GAL80-mediated repression of GAL4 (38) in place in W303 cells. On the other hand, domains derived from the herpes simplex virus TF VP16 have shown limitations before when adopted in yeast (39). In contrast, SYRAP and UbSYRAP activated *HRPE-NLUC* and *HRPE*_*ADH*_*-NLUC*, causing 4-to 6-fold NLUC enhancement over their GAL4STE12 control (**Fig. 3A**). A similar trend was also visible with *HRPE*_*Ω*_*–NLUC*, although masked by the variability of this highly active promoter. A preliminary test of the oxygen sensitivity of the two SYRAP versions was made in Arabidopsis, where the Cys-NDP regulation exists natively. Transient transformation of plants expressing a stable *HRPE*_*ADH*_*-NLUC* reporter (40) showed that both TFs retained O_2_-dependent regulation (**Fig. 3B**). Their strong constitutive expression boosted reporter activity far beyond the levels allowed by the native ERFVII machinery.

Finally, we implemented the O_2_-dependent regulation in the yeast circuit with a plant-derived PCO module. This modification is in fact known to enable a synthetic Cys-NDP in *S. cerevisiae* (27). Preliminarily, we made sure that a nuclear-localized PCO construct was able to process the model substrate DLOR (27) (**SI Appendix, Fig. S3B**). We then opted to assess the performance of the circuit in different ranges of output activation; based on the previous results, we thus tested SYRAP and UbSYRAP susceptibility to PCO when combined to *HRPE-* or *HRPE*_*ADH*_*-NLUC* modules (**Fig. 3C**). Both PCO and normoxia were required to prevent the activation of the promoters, but the regulation was partially masked in the high output range associated with *HRPE-NLUC*. The best regulatory range was attained by combination of UbSYRAP with *HRPE*_*ADH*_*-NLUC*, where low output was associated with effective PCO- and O_2_-dependent degradation of the TF (**Fig. 3D, E** and **SI Appendix, Fig. S3C)**. We thereby demonstrated that a minimal set of plant-derived modules is sufficient to install O_2_-dependent transcriptional circuits in *S. cerevisiae*.

Relying on the dual DNA binding capacity of SYRAP effectors, we explored the possibility to associate this synthetic regulation with *UAS*_*GAL*_ promoters (**Fig. 3F**). To this end, we deployed two *UAS*_*GAL*_ genomic insertions, P*HIS3UAS*_*GAL1*_*-HIS3* and P*GAL1-LACZ*, carried by the MaV203 strain (41). Aerobic growth of cultures expressing either SYRAP or UbSYRAP on selective histidine-deficient media was indeed PCO-dependent (**SI Appendix, Fig. S4A-B**). The β-galactosidase assay confirmed strong aerobic activation of *LACZ* by the two SYRAPs, activation prevented if PCO was expressed (**SI Appendix, Fig. S4C**). Also on solid media, SYRAP and UbSYRAP colonies showed PCO-mediated growth repression in aerobic conditions (**SI Appendix, Fig. S4D**). As expected, PCO-positive UbSYRAP colonies developing in hypoxia were instead undistinguishable from PCO-negative colonies (**Fig. 3G**). These experiments suggest that ERFVII-based transcriptional circuits represent a viable strategy to achieve O_2_-dependent control of complex responses, such as growth, in yeast without interference with the endogenous O_2_ signalling pathways (*i*.*e*., orthogonally).

### Fast and reversible responses to O_2_ variations are produced by the minimal ERFVII-based transcriptional circuit

The assembly of PCO, UbSYRAP and *HRPE*_*ADH*_*-NLUC* modules was adopted for subsequent experiments and designated as MynOx, for Minimal yeast Oxygen-responsive circuit (**Fig. 3F**). MynOx yeast was meant to serve as a heterologous model for plant hypoxic regulation; a precondition to its use was thus to ensure that yeast cells could be exposed to low oxygen in a way that faithfully approximated the conditions experienced by seedlings in **Fig. 1A**.

We initially monitored MynOx output in hypoxic liquid cultures, while recording O_2_ concentration in the medium with a sensor spot (as described in (42)). O_2_ depletion was associated with progressive output increase (**Fig. 4A**), as expected only in presence of PCO. The treatment was protracted until the cultures reached complete anoxia, after which they were re-oxygenated. By the time dissolved O_2_ was restored to pre-hypoxic levels, the NLUC output had fallen from 4-fold induction to 2-fold the initial aerobic values (**Fig. 4A**), indicating that MynOx generates quantitative and reversible responses to O_2_ fluctuations. However, the culture medium underwent slow equilibration with the hypoxic atmosphere, which resulted in delayed output. Moreover, failure to maintain a steady-state level of dissolved O_2_, likely due to fast consumption by proliferating cells, precluded the possibility to titrate MynOx output as a function of O_2_ concentration. To overcome these limitations, we applied a different treatment set-up, re-suspending the cultures in media that had been pre-equilibrated under hypoxia. This time, stationary O_2_ levels could be maintained for at least 1 h, enabling a time-resolved analysis of the response (or “hypoxic chase”, **Fig. 4B**). Cultures equilibrated at 1% O_2_ quickly reached a plateau of NLUC signal in 10 minutes, whereas no response was observed in cultures exposed to very mild hypoxia (10% O_2_) (**Fig. 4B**). We concluded that MynOx could react promptly to acute hypoxia, depending on the perceived level of oxygen in cells.

**Figure 4.**
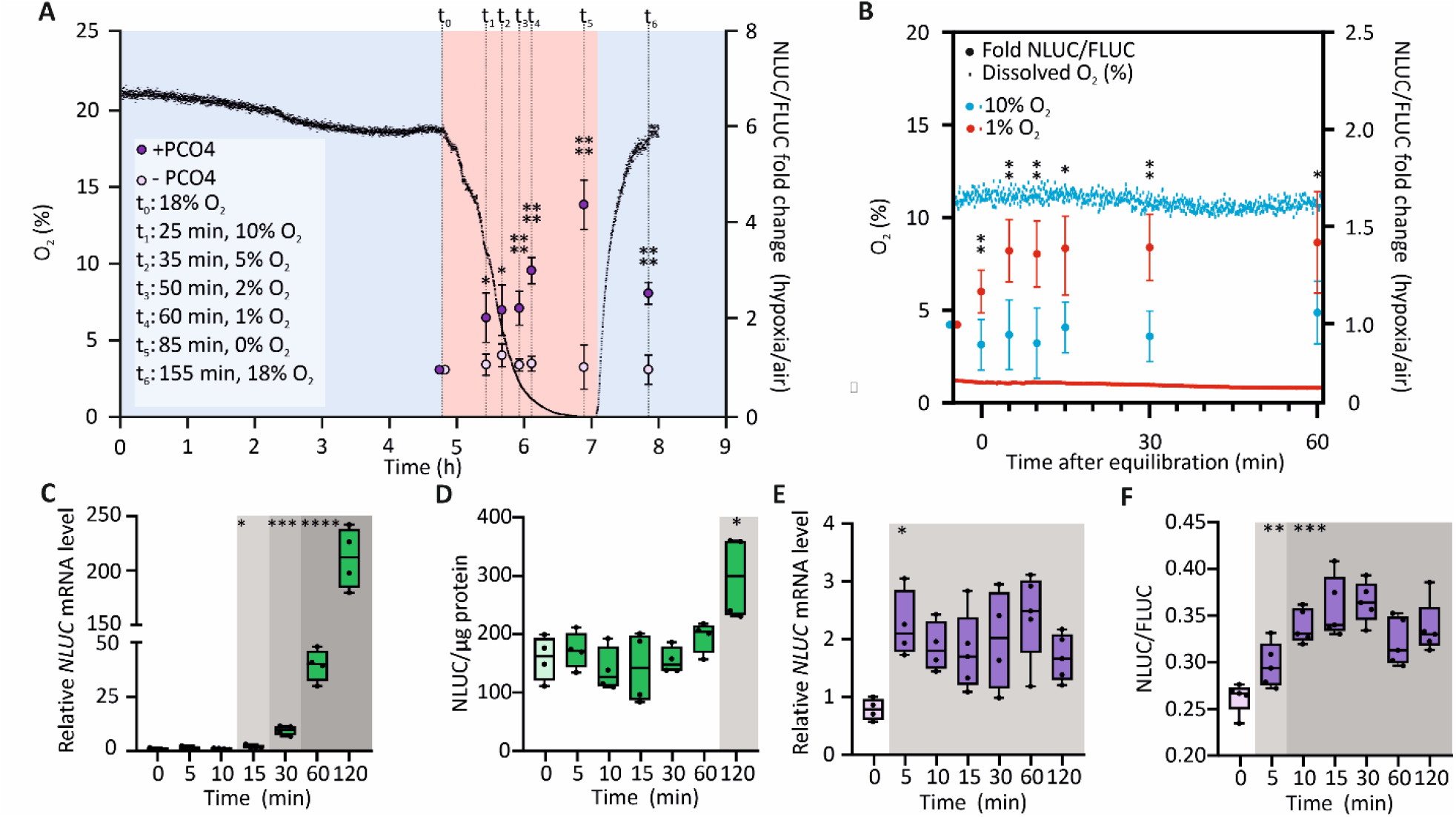
Recapitulation of plant hypoxic dynamics in yeast. **(A)** Evolution of *HRPE*_*ADH*_*-NLUC* output in cells expressing PCO4 (purple dots) or GUS (yellow dots). Cultures grown for 5 h in normoxia (t_0_ of the treatment) were maintained in a 1% O_2_ hypoxic atmosphere (red shading). Re-oxygenation (blue shading) was applied after anoxia was reached in a reference culture, where dissolved O_2_ levels were monitored (black profile), and was protracted until O_2_ reached back the t_0_ level (18%, v/v). Data are expressed as fold changes of NLUC/FLUC activity between hypoxic and normoxic (t_0_) samples (mean ± SD, n=5). **(B)** Output of MynOx cultures in “hypoxic chase” experiments. Cultures grown for 5 h in normoxia were resuspended in media equilibrated at 10% (blue) or 1% O_2_ (red) and incubated under the corresponding hypoxic atmosphere. NLUC activity (dots) was monitored at 5, 10, 15, 20, 30 and 60 min after culture equilibration with the hypoxic media and expressed as fold change of NLUC/FLUC activity over normoxia (mean ± SD, n=5). Coloured profiles show O_2_ concentration (% v/v) in the culture. **(C)** *NLUC* expression in 7-day-old *HRPE*_*ADH*_*:NLUC* seedlings from vertical plates, normalized to the *UBQ10* housekeeping gene and made relative to a t_0_ sample (n=4). **(D)** Relative NLUC activity in the same seedlings. **(E)** *NLUC* expression in MynOx thin colonies, normalized to *Actin 1* and made relative to a t_0_ sample (n=4-5). **(F)** NLUC/FLUC activity in the same colonies. Asterisks indicate significant differences (*, 0.01≤p<0.05, **, 0.001≤p<0.01, ***, 0.0001≤p<0.001, ****, p<0.0001) after Student’s t-test on pairwise comparisons between + and −PCO4 (A), 1% and 10% O_2_ (B), or after multiple t-tests between each timepoint and t_0_ (normoxia, C-F).

These observations directed us to shift from liquid cultures to thin-layered colonies, whose morphology should prevent the formation of oxygen gradients. Cells in such colonies were expected to remain equilibrated with the atmosphere for longer time than cultures and better approximate the speculated situation in seedling tissues. We thus compared the dynamic output of MynOx colonies with *HRPE*_*ADH*_*-NLUC* Arabidopsis seedlings, over a timing of hypoxia. *NLUC* mRNA built up quickly both in plants (**Fig. 4C** and **SI Appendix, Fig. S5A**) and yeast (**Fig. 4E**). Remarkably, the rate of *NLUC* accumulation was the same as the *ADH1* transcript (**SI Appendix, Fig. S5A** and **S5C**), suggesting that the synthetic HRPE promoter could recapitulate the dynamics of endogenous genes and pointing at a decisive role for the 5’-UTR in *ADH1* regulation. The NLUC output showed sizeable delay in plants (**Fig. 4D** and **SI Appendix Fig. S5B**), whereas it closely followed mRNA pattern in yeast (**Fig. 4F**). The lag in plants may be explained due to selective protein translation (43), in contrast with hypoxic promotion of protein synthesis in yeast (44, 45).

After the initial induction, nonetheless, the responses diverged with time. *NLUC* mRNA kept accumulating until the end of a 2 hour-long treatment in seedlings (**Fig. 4C**), while in yeast it reached a steady-state within few minutes (**Fig. 4E**). Moreover, the response range was incomparably smaller in MynOx than Arabidopsis (2.5-fold *NLUC* induction in yeast, 200-fold in seedlings). These data highlight that the minimal circuit, although able to trigger *HRPE*_*ADH*_*-NLUC* expression, was insufficient to establish a complete transcriptional response in yeast.

### Sustained induction of anaerobic gene expression requires positive ERFVII-regulated feedback

A major difference between the two *HRPE-*inducing devices compared above resided in the fact that plants incorporate a hypoxia- and ERFVII-regulated feedback loop. In Arabidopsis, constitutively expressed ERFVII paralogs (AtRAP2.2/3/12) are initially stabilized under low oxygen, upon inhibition of constitutively expressed PCOs (AtPCO3/4/5), and in turn trigger the expression of hypoxia-inducible *ERFVII* (*AtHRE1/2*) and *PCO* (*AtPCO1/2*) genes (7, 46). To estimate the impact of this feedback in the plant hypoxic response, we introduced it into MynOx. We transformed *MATα* W303 cells with two ERFVII-inducible modules, HRPE_ADH_-UbSYRAP (U) and HRPE_ADH_-PCO1 (P), or their respective HRPE_ADH_-GUS control constructs (G). Mating of each *MATα* strain with MynOx (*MATa*) wired the inducible modules to the existing minimal circuit (**Fig. 5A**). The four diploid strains obtained, designated by their *MATα*-specific genotype hereafter, contained no new inducible module (GG), HRPE_ADH_-UbSYRAP only (UG), HRPE_ADH_-PCO1 only (GP), or both HRPE_ADH_-UbSYRAP and HRPE_ADH_-PCO1 (UP). After 6 h hypoxia, control diploid cultures (GG) reached comparable output to the parental haploid strain (**Fig. 5B, Fig. 2C**), whereas the hypoxic output expanded significantly by addition of the HRPE_ADH_-UbSYRAP module (UP and UG) (**Fig. 5B**). A chimeric factor derived from AtHRE2 (HRPE_ADH_-UbSYHRE), unable to bind the HRPE element directly (30), left the minimal output unchanged (**SI Appendix, Fig. S6A-B**). Finally, addition of HRPE_ADH_-PCO1 alone (GP) produced negligible effects (**Fig. 5B**).

**Figure 5.**
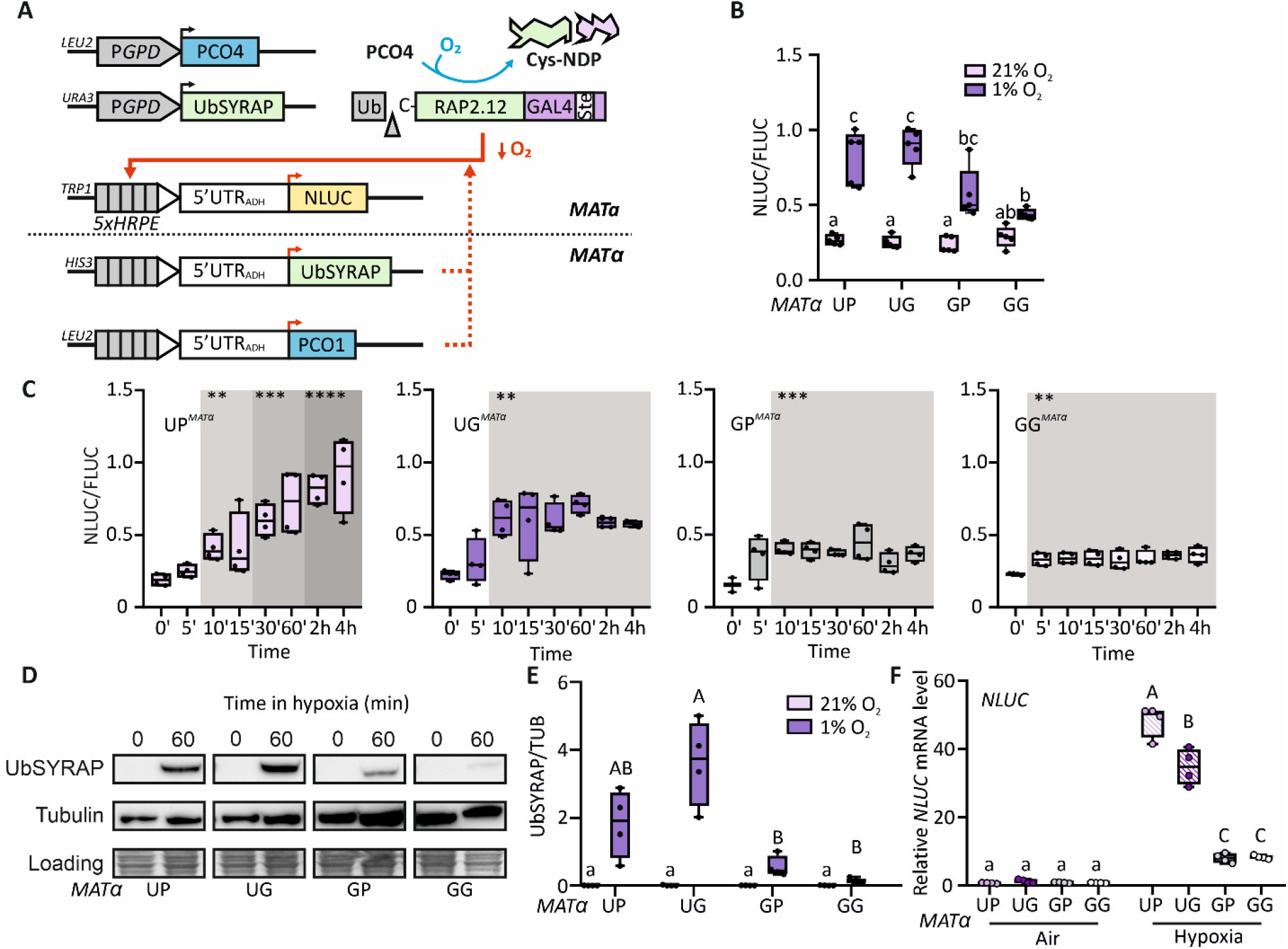
Introduction of a positive hypoxia-responsive feedback circuit in yeast. **(A)** Schematics of the modules expressed in the diploid W303 strain. The ERFVII-inducible modules HRPE_ADH_-UbSYRAP and HRPE_ADH_-PCO1 were introduced in *MATα* cells, to be combined with the constitutive modules from MynOx^*MATa*^, controlled by glyceraldehyde 3-phosphate promoters (P*GPD*). Hypoxia triggers positive feedback through activation of the inducible modules. **(B)** Relative NLUC activity (NLUC/FLUC; n=5) in diploid cultures after 6 h hypoxic (red boxes) or aerobic treatment (blue boxes). The *MATα* genotypes mated with MynOx^*MATa*^ consisted of combinations of HRPE_ADH_-UbSYRAP (U), HRPE_ADH_-PCO1 (P) or HRPE_ADH_-GUS (G, negative control) modules, integrated either at the *HIS3* or *LEU2* locus. **(C)** Evolution of NLUC activity (n=4) in diploid colonies moved from normoxia (t_0_) to 4 hour-long hypoxia (1% O_2_). **(D)** UbSYRAP protein abundance in normoxic diploid cultures (t_0_) or after 1 h incubation in hypoxia in pre-equilibrated medium. **(E)** Normalized band intensity to α-tubulin, in independent biological replicates (n=4). **(F)** *NLUC* mRNA abundance, normalized on *ACT1* expression, in different diploid cultures treated as in (D). Data are relative to an aerobic GP^*MATα*^ sample. Asterisks and shadings indicate significant differences (**, 0.001≤p<0.01, ***, 0.0001≤p<0.001, ****, p<0.0001) after multiple Student’s t-tests between each time point in hypoxia and t_0_. Different letters in (E) and (F) indicate statistically significant differences among normoxic (lowercase) or hypoxic samples (uppercase), after One-way Anova, while in (B), they indicate differences between groups and treatments after Two-way Anova was performed (p<0.05).

We then profiled the response of diploid genotypes to steady-state hypoxia in a time-resolved experiment (**Fig. 5C**). The induction in the absence of feedback was fast but contained (GG), with similar range to the haploid MynOx (**Fig. 3F**). Introduction of the inducible PCO module (GP) did not alter circuit behaviour. The inducible TF alone (UG) increased NLUC induction to 3-fold but could not prolong the response beyond 10 min treatment. With the full feedback (UP), instead, the initial induction was maintained and the time of response stretched, at a non-linear rate, up to 2 hours into the treatment (**Fig. 5C**). As a result, UP enhanced the 1.5-fold dynamic range of GG at 4 h to 5-fold (**Fig. 5C**). The UbSYRAP protein accumulated to higher levels in UG and UP under hypoxia than in other genotypes (**Fig. 5D-E** and **SI Appendix, Fig. S6C-D** and **S7**), supported by higher *UbSYRAP* expression **(SI Appendix, Fig. S6E)**, and *NLUC* expression increased by 4-folds in steady-state conditions (**Fig. 5F**). Early hypoxia (5’) induced *NLUC* in all genotypes except for GP (**SI Appendix, Fig. S8**), but *NLUC* expression could be maintained over a 4-hour time-course only when the inducible UbSYRAP module was present, in UP and UG, while the expression declined in GG and GP. The differences among genotypes were not associated with general alterations in endogenous transcription, as shown by use of yeast anaerobic gene markers (**SI Appendix, Fig. S6F**), indicating that the synthetic circuit acted orthogonally to yeast anaerobic pathways.

By implementing basic ERFVII-regulated feedback in our simplified circuit, we demonstrated that combination of a hypoxia-inducible ERFVII module with an inducible PCO module was sufficient to expand the time of response and dynamic range of the output. Altogether, the yeast model indicates that constitutively expressed RAP2-type *ERFVII*s are responsible and sufficient for the immediate activation of target genes, but a low oxygen-inducible ERFVII/PCO feedback mechanism is required to prevent a fast decline of the response and support higher induction rates.

### Mathematical modelling reveals key circuit features to achieve fast transcriptional responses to hypoxia

We constructed four mathematical models, backed up by the output data collected, to interpret the behaviour of the diploid strains. The models ranged in complexity from the essential circuit based on a single ERFVII and a PCO (GG), to the simplified feedback loop including hypoxia-inducible copies of both modules **(Fig. 6A)**. The coupled Ordinary Differential Equations (ODEs) defining every model (**SI Appendix, Table S2**) were built using the reaction rates reported in **SI Appendix, Table S3**. Except for the kinetic variables for PCO4 and PCO1, which were re-fitted from *in vitro* data (12) (**SI Appendix, Fig. S9A**), and the rates of UbSYRAP synthesis and degradation (k1 and k2), calculated from (27) (**SI Appendix, Fig. S9B**), most parameters were built to best fit our experimental data (**SI Appendix, Table S4**). The obtained simulations compared well with the experimental NLUC induction (NLUC/FLUC; **Fig. 5C, Fig. 6B**) indicating that the nonlinear fitting was accurate, and the parameters reasonable (**SI Appendix, Table 4**). Additionally, we could also simulate UbSYRAP dynamics, which we could otherwise only estimate from immunoblotting (**SI Appendix, Fig. S6C-D**). Compatible with the experimental observations, UbSYRAP amount decreased over time in GP, but not in UP (**Fig. 6B**), suggesting that the hypoxia-inducible PCO retains activity under 1% O_2_ atmosphere, but repression is overcome by the presence of a hypoxia-inducible copy of the TF.

**Figure 6.**
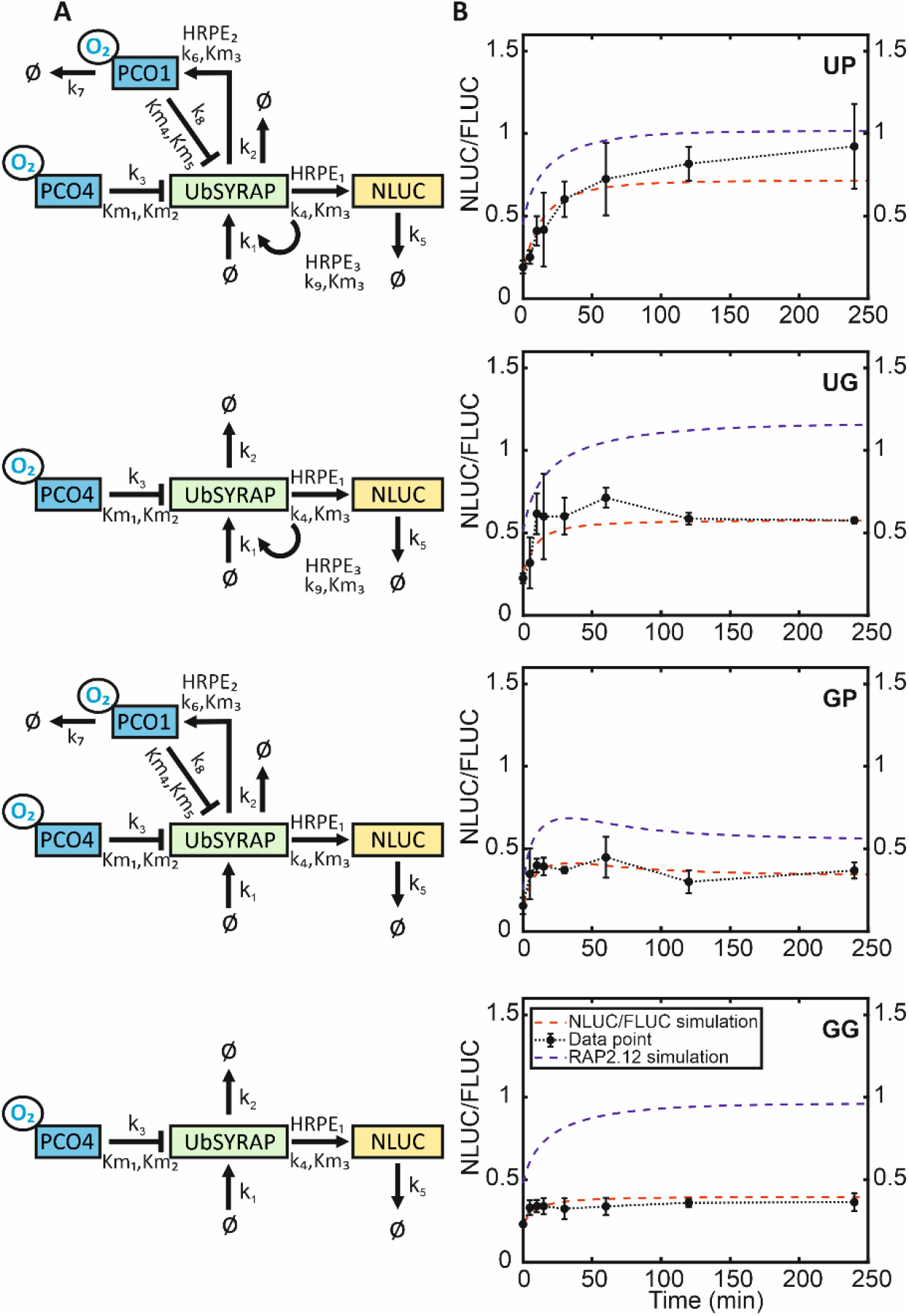
Mathematical modelling of the synthetic O_2_-responsive circuit of yeast. **(A)** Schematic architecture of the models corresponding to the four genetic circuits illustrated in **Figure 5**. All species included in the models (UbSYRAP, PCO1/4, O_2_ and NLUC, output corresponding to the NLUC/FLUC output in yeast) are framed, while the ‘Ø’ symbol stands for ‘null’. **(B)** Computational simulation of NLUC activity based on the previous models (red line), overlapped with the experimental data from **Fig. 5C** (NLUC/FLUC, black line). The adjusted UbSYRAP concentration (blue line, “RAP2.12 simulation”) was obtained from the steady-state simulations, fitted as described in SI Appendix, Extended methods from (27).

We observed changes in the parameter values across the four models, revealing that the binding rate of RAP2.12 to different HRPE promoters significantly influences system behaviour. Notably, the binding rates of RAP2.12 to *HRPE*_*ADH*_*-NLUC* (HRPE_1_, output module) varied among GG, GP, UG and UP models (0.07, 0.06, 0.04, and 0.02, respectively; **SI Appendix, Table S4**). Introduction of a hypoxia-inducible RAP2.12 copy (*HRPE*_*ADH*_*-UbSYRAP*) in UG and UP showed a more substantial HRPE_1_ reduction than *HRPE*_*ADH*_*-PCO1* only (GP). Similarly, the predicted binding rates of RAP2.12 to *HRPE*_*ADH*_*-PCO1* (HRPE_2_) and *HRPE*_*ADH*_*-UbSYRAP* (HRPE_3_) were lower in UP, with both inducible components present. The lowest HRPE_1/2/3_ values in UP (**SI Appendix, Table S4**) suggest a competitive interaction among promoters for RAP2.12 recruitment, leading to further modulation in their individual binding efficiencies.

We examined how the full-feedback model UP, which resembles plant cell regulation most closely, responded to variations of individual parameters (**SI Appendix, Fig. 10A**), based on NLUC response time (t_90%_-t_10%_, a proxy of NLUC/FLUC induction speed) as model performance metric. Interestingly, among the top-ranking parameters influencing the response time, five were linked to PCO1: k_6-8_ (PCO1 degradation, PCO1 synthesis and RAP2.12 oxidation rate by PCO1) and Km_4-5_, accounting for the re-fitted Michelis-Menten constants for O_2_ and RAP2.12 as PCO1 substrates (**SI Appendix, Fig. 10B-C and Table 4**). Overall, high PCO1 activity under hypoxia resulted to be associated with shorter response time. Our analysis puts forward a potential strategy to enhance the speed of plant hypoxic responses through PCO1 engineering, leading to higher affinity for O_2_ and RAP2.12 (lower Km_4-5_), elevated catalytic rate for RAP2.12 oxidation (k_8_) or higher PCO1 abundance (higher k_6_, or lower k_7_) (**SI Appendix, Fig. 10C**).

## Discussion

Plant ERFVII transcription factors have long been identified as responsible to orchestrate the expression of low oxygen-inducible genes in plants (24, 47). The ERFVII proteins are normoxically degraded via the cysteine-(or PCO-) branch of the N-degron pathway (Cys-NDP) and stabilized when oxygen levels fall (9, 10), which creates a mechanistic link between the ERFVII and the observed responses. However, the expected prompt activation of hypoxia-responsive genes seems to conflict with evidence of ERFVII protein stabilisation after longer treatments, collected using fluorescent reporters and immunoblots (18, 33). This has led to speculation that additional mechanisms are needed to generate the first response to hypoxia in plants, potentially acting upstream of the ERFVIIs, or in parallel to them but with faster dynamics (23, 48). Transcriptomic studies have defined the signature of acute hypoxia as early as 30’ into the treatment (29, 49), nonetheless quicker induction of transcription has been found with marker genes (30, 50). Here, we showed that anaerobic markers (HRGs) are already upregulated after 5 minutes exposure of plant tissues to hypoxia (**Fig. 1A**). Importantly, we obtained evidence that correlate the timing of the transcriptional response with increased RAP2.3^3xHA^ protein amount (**Fig. 2C-D**), in support of the hypothesis that signalling events leading to fast ERFVII stabilization are a requisite for HRG induction.

Anaerobic transcription in seedlings was as expected sustained, until at least 4 h treatment (**Fig. 1A** and **C, SI Appendix, Fig. S5C**). Hypoxia causes coordinated epigenetic modifications in up-regulated HRG regions. The chromatin at these loci becomes more accessible early under low oxygen, the repressive H2A.Z histone form is displaced, and transcription is promoted by the rise of activating histone modifications, such as H3K9ac (34, 51). Chromatin remodelling factors are also recruited onto HRG loci through the Mediator complex, after ERFVII binding to their upstream sequences (52–54). In agreement with the role of ERFVIIs in promoting the accessibility of up-regulated genes, we found a correlation between the different profiles of HRG induction and the distribution of HRPE *cis*-motifs bound by RAP-type ERFVIIs (**Fig. 1B**). This suggests that ERFVIIs can determine both the amplitude and speed of the hypoxic responses. The synthetic *HRPE*_*ADH*_ promoter showed intermediate response time in plants (like the endogenous gene *ADH1*, **SI Appendix, Fig. S5A**), despite five tandem repetitions, implying that the exact profile of HRG expression depends on HRPE arrangement. The same promoter construct was quickly induced by hypoxia in yeast but could not reach the induction levels observed in plants (**Fig. 5F** and **SI Appendix, Fig. S8**). Despite conservation of the core chromatin remodelling machinery in yeast (55, 56), inefficient association with ERFVII transcriptional complexes may partially account for these discrepancies in *NLUC* output.

Our main goal was to measure the contribution of ERFVII conditional proteostasis to HRG induction, disentangling it from other concurrent signalling events associated with hypoxia in plant cells. Observing the genuine effects of the Cys-NDP on ERFVII activity entails the ability to insulate that pathway from others potentially impinging on the same protein targets. Synthetic biology provides the ideal framework to manage the reduction of complex biological behaviours into simpler functioning schemes (26, 57). Abstraction of signalling elements from the native plant host has been successfully attempted before, facilitated by the intrinsic modularity of signalling mechanisms (58, 59). For instance, engineering of core components of plant nuclear auxin signalling in yeast has unveiled the hierarchical relationships among Aux/IAA factors (60), otherwise masked by genetic redundancy of the pathway, feedback and cross-regulation. We pursued our goal with a similar strategy, reconstituting an O_2_-dependent transcriptional circuit by abstraction of essential modules of the plant Cys-NDP.

The minimal O_2_-responsive circuit (MynOx, **Fig. 3F**) demonstrated that coupling of an ERFVII-based effector with a PCO sensor was enough to provide hypoxic activation of *HRPE*_*ADH*._ The reliability of a heterologous model of plant oxygen sensing also depends crucially on the possibility to control O_2_ supply to cells. We identified thin-layered colonies as the best material to approximate the effects of plant seedling tissues exposure to hypoxia (**Fig. 4**). The comparable speed of output activation in yeast colonies or Arabidopsis seedlings exposed to the same hypoxic atmosphere (**Fig. 4C, E** and **SI Appendix, Fig. S5**), indicates that the most immediate response in Arabidopsis relies exclusively on PCO-dependent signalling. Recently, fast translocation of the calcium-dependent protein kinase CIPK12 to the nucleus has been suggested to protect the ERFVII factors from degradation during hypoxia in Arabidopsis, potentially providing a stabilization mechanism independent of the Cys-NDP (20). The CIPK12 pathway is unlikely to be conserved in yeast, where hypoxia alone has not been shown to directly trigger calcium signalling (61) Although we cannot exclude phosphorylation of the synthetic RAP2.12, hypoxia could only induce MynOx output (or a growth output) in cells expressing PCO (**Fig. 4A** and **3G**).

The design of MynOx highlighted the importance to adapt the effector module to yeast transcriptional requirements. The plant-native activation domain of RAP2.12 was not functional in yeast (**Fig. 3A** and **SI Appendix, Fig. S4**), nor was the synthetic VP64 domain (**Fig. 3A**). In both cases, specific mediator complex subunits, absent in *S. cerevisiae*, might be required to recruit the RNA polymerase (59, 62). Cognate HRPE promoters were taken from Panicucci et al. (2020), with no further adaptation. All sequences contained a −47 to +36 fragment from the 35S promoter of Cauliflower Mosaic virus (63), transcriptionally functional in yeast and presenting a compatible TATA box consensus sequence (64, 65) 31 nt upstream of the TSS. Addition of the omega leader sequence of Tobacco Mosaic Virus (66) caused intense NLUC translation in basal conditions and severely limited the dynamic range of the output; in contrast, the 5’-UTR sequence from Arabidopsis *ADH1* conferred broad dynamic range to MynOx (**Fig. 3A, C** and **SI Appendix, Fig. S3A**). These results highlight the importance of the correct choice of 5’UTR regions for the precision control of yeast genetic circuits (67, 68).

Plant *ERFVII* and *PCO* gene families include hypoxia-inducible members (47, 69). The divergence of hypoxia-inducible (or B-type) *PCOs* is puzzling, but their conservation in Spermatophytes (46) indicates that they are under selective pressure. The limited range of response displayed by MynOx, in contrast with the output of *HRPE*_*ADH*_ seedlings (**Fig. 4C-F** and **SI Appendix, Fig. S5A-B**), prompts to speculate that the hypoxia-inducible components of the regulatory mechanism are instrumental to the amplitude of *HRG* response. MynOx expansion by rational design (**Fig. 5A**) in fact improved the output only in presence of both *HRPE*_*ADH*_-driven modules (**Fig. 5C, 5F** and **SI Appendix, Fig. S8**). B-type PCOs have been proposed to optimize ERFVII processing during O_2_ fluctuations, however their functions during acute hypoxia could not be disentangled through genetics so far (46). Our data show that B-type PCOs are specifically required to regulate the amplitude and duration of the anaerobic response. PCO1 introduction in the circuit produced sizeable effects in 1% O_2_ atmosphere, hinting at different kinetic properties of these enzymes *in vivo* than measured *in vitro* (12). The fitted model predicted a progressive depletion of the constitutive RAP2.12 pool in GP under hypoxia, resulting in low output (**Fig. 5B**) and low target gene expression (**SI Appendix, Fig. S8**). This outcome matches well with the early decline in *HRG* transcripts observed in *hre1hre2* mutants (47). Here, lack of inducible *ERFVIIs* able to counteract the B-type PCOs led to plant inability to sustain the anaerobic response. Our heterologous model thus suggests that the co-evolution of hypoxia-inducible *PCO* and *ERFVII* genes installed a regulatory loop that shapes plant transcription under stress. The parameter sensitivity analysis further detailed the role for B-type PCOs in hypoxia, specifically at tuning the output speed (**SI Appendix, Fig. S10**). This provides further insight into rational B-type PCO modifications increasing their concentration or affinity towards O_2_ and RAP2s, in order to speed up transcriptional responses to hypoxia in plants.

The modular architecture of our heterologous model also enabled us to pinpoint the competition among HRPE-containing promoters during hypoxia. The four fitted models showed that RAP2.12 binding rate to the promoter of the output module (HRPE_1_) was influenced by the presence of other RAP2.12-inducible modules, assuming its lowest value in the UP model (**SI Appendix, Table S4**). This highlights the critical role of ERFVII binding kinetics in modulating HRG expression dynamics. According to competition theory models, multiple promoters sharing the same *cis*-elements are expected to compete for the recruitment of cognate TFs present in limited pools (70, 71). The theory has found broad implications in molecular contexts (71–73). In the case of co-expression networks coordinated by individual TF families, such as the plant HRG network, competition (also indicated as “TF titration”) can explain the diversification of target gene expression patterns (71). According to the regulatory model, titration phenomena can generate non-linear responses, such as digital outputs as in the winner-take-all (WTA) effect (74), or define the expression hierarchy of the targets (70, 75). The competition phenomenon revealed by our model provides a possible explanation for the expression kinetics observed in **Fig. 1**, with fast HRG induction being dictated by stronger HRPE arrangements in presence of limited RAP transcription factor pools (compatible with the speculative model suggested by Burger 2010).

By virtue of its modularity, our synthetic yeast model represents an ideal tool to compare the binding affinity of different ERFVIIs to HRPEs and thereby start disentangling the questions related to the functional specificities of RAP2-type paralogs in plants (76, 77), which have remained unaddressed so far. We could confirm that HRE-type ERFVII are unable to associate with HRPE promoter elements (**SI Appendix, Fig. S6A**), as reported before by Gasch et al (2016) and Lee & Bailey-Serres (2019) (30, 34). This evidence indicates that re-design of the output module by incorporation of GCC-rich motifs (34) should make it possible to implement HRE-based hypoxic feedback in the synthetic model and thereby quantify the share of HRE regulation in the long-term anaerobic output.

In conclusion, in this study we engineered a “plantified” yeast strain to serve as foundational model of synthetic oxygen-dependent transcriptional devices. We demonstrated that this is a viable strategy to extrapolate quantitative information on the impact of the Cys-N degron pathway on hypoxic gene expression that would be difficult to extract from the native plant host, using genetic approaches. Moreover, the minimal MynOx circuit constitutes a unique orthogonal switch controlled by hypoxia that may be further exploited to introduce oxygen-dependent responses in yeast.

## Materials and methods

### Plant material

Col-0 *Arabidopsis thaliana* was used as wild-type. The pentuple *erfvii* mutant (31) and the overexpressing line *35S:RAP2*.*3*^*3xHA*^ (33) were kindly provided by Mike Holdsworth. The *HRPE*_*ADH*_*:NLUC* reporter line was described by (40). Seven day-old Arabidopsis seedlings were subjected to hypoxic or chemical treatments. Plants were grown on 2.15 g L^-1^ MS medium (Duchefa) supplemented with 10 g L^-1^ sucrose, either in 6-well plates, or vertical plates in presence of 9 g L^-1^ agar, as described previously in (78). Transient expression of SYRAP and UbSYRAP constructs was accomplished by agroinfiltration of three week-old *HRPE*_*ADH*_*:NLUC* plants, according to (37). Three days after infiltration, leaves were excised and incubated in 1 mL of water under normoxic or hypoxic conditions, with 150 rpm shaking and 23°C in the dark.

### Yeast strains, culture and transformation

The haploid *S. cerevisiae* strains MaV203 (Thermo-Fisher Scientific #11445012), W303 1A (*MATa*) and W303 1B (*MATα*) (Scientific Research and Development, GmbH, #20000A and #20000B) were used. Fresh cells from YPDA plates were transformed following the PEG/LiAc/SS DNA method (79). A detailed transformation protocol is described in **SI Appendix**, Extended methods. All strains generated in this study are listed in **SI Appendix, Table S5**. Unless differently specified, yeast was grown on synthetic yeast dropout (SD) medium, with appropriate supplements, as described in (27). Spots were prepared from exponential cultures (OD_600_=0.75). Approximately 10^5^ cells were spotted on solid SD medium (-his-trp-leu-ura) and grown for 28 h at 30°C prior to treatments. For gene expression and luciferase measurements, cells were sampled by gently scratching the surface of colonies, avoiding contaminations with the agar, resuspended in a tube and flash frozen.

### Cloning of yeast and plant components

Entry vectors were generated by Gateway BP Clonase reaction between pDONR201 (Thermo Fisher Scientific) and attB containing inserts. Strings were synthesized through GeneArt® (Thermo-Fisher Scientific), **SI Appendix, Table S6**. Cloning procedures and sequence information of the inserts can be found in SI Appendix, Extended methods, and the specific primers used in **SI Appendix, Table S7**. Entry plasmids (**SI Appendix, Table S8**) were recombined into suitable destination vectors using the Gateway− LR Clonase− II Enzyme mix (Thermo-Fisher Scientific). Combinations of yeast expression vectors were transformed into the different yeast strains as detailed in **SI Appendix, Table S6**.

### Low oxygen treatments

Treatments were carried out in a sealed hypoxic glovebox with temperature control (Coy Lab Products). Arabidopsis seedlings and leaves were kept in the dark at 23°C, under 120 rpm continuous shaking, for the duration of the hypoxic treatment. The plant material was sampled inside the glovebox by flash freezing in liquid nitrogen.

Hypoxia was applied to diluted yeast suspensions dispensed in 200 µl volume in 96-well plates (microcultures), or to 5- or 10-ml cultures in individual vials, as specified for the different experiments. The treatments consisted in the incubation under 1% O_2_ atmosphere (v/v), at 30 °C and with 150 rpm shaking. Culture production and handling has been described before (78). In “hypoxic chase” experiments, media were pre-equilibrated with the hypoxic atmosphere in the glovebox, set at the desired O_2_ concentration (10% or 1% O_2_ v/v). Five mL cultures diluted to OD_600_=0.1 were first pre-grown for 5 h in aerated conditions to exponential phase, then recovered by centrifugation (5000 g, 2 min) and resuspended in the same volume of pre-equilibrated media directly inside the hypoxic glovebox. For the duration of the treatment, O_2_ concentration in the media was measured with a FireSting®-GO2 meter (PyroScience), thanks to a fluorescent oxygen sensor spot (OXSP5, PyroScience) attached inside the container with the medium. Dissolved O_2_ was also monitored in the same way inside a separate culture for the duration of the treatment, to ensure the maintenance of steady-state O_2_ levels across all sampling points (42).

### Reporter assays

Luciferase activity was assessed with the aid of commercial kits, following the manufacturer’s recommendations. Approximately 5×10^5^ yeast cells were harvested from growing cultures or spots, flash frozen in N_2_ and resuspended in 150 μl Passive Lysis Buffer (PLB, Promega). NLUC activity was measured and normalized to FLUC activity (NLUC/FLUC) using the Nano-Glo® Dual Luciferase Assay System (Promega). In the case of Arabidopsis samples, 15 seedlings or one agroinfiltrated leaf were frozen, ground to fine powder with mortar and pestle and suspended in 1 mL PLB. NLUC signal was measured with the Nano-Glo® Luciferase Assay System (Promega) and normalized to the amount of total proteins, quantified using the Pierce− BCA Protein Assay Kit (Thermo-Fisher Scientific). DLOR activity was measured as described in (27) using the Dual-Luciferase® Reporter Assay System (Promega). Detailed information on the UAS_GAL_ activation experiments performed in MaV203 strains is provided in SI Appendix, Extended methods.

### Gene expression analyses

Total RNA was isolated from Arabidopsis samples as described before (18) and from yeast samples as described in (78). One μg total RNA was processed to cDNA with the Maxima cDNA synthesis kit (Thermo-Fisher Scientific), according to the manufacturer’s protocol. Real-time qPCR was performed with 12.5 ng cDNA template, using the PowerUp SYBR® Green Master Mix (Thermo-Fisher Scientific) and a CFX384 detection system (Bio-Rad). Gene-specific qPCR primers are listed in **SI Appendix, Table S9**. Gene expression was expressed as relative to the yeast housekeeping gene *ACT1* (YFL039C) and the Arabidopsis housekeeping *UBQ10* (At4g05320), through the comparative ΔΔCt method (80).

### Immunoblotting

Detailed information on protein extraction from plant and yeast material, immunoblotting and detection is provided in SI Appendix, Extended methods.

### Promoter analysis

The intergenic regions upstream of ATG of the nine selected HRGs (*ADH1, HRA1, PDC1, PGB1, PCO1, LBD41, HRE2, SAD6* and *SUS4*) were extracted using Integrated Genome Browser® (v. 10.0). The individual motif occurrence tool FIMO (https://meme-suite.org/meme/tools/fimo) was used to find Hypoxia Response Promoter Elements (HRPEs) from the available consensus sequences (Gasch et al., 2016) with a filtering p-value of 0.001 for positive hits.

### Mathematical modelling of NLUC, RAP2.12 and PCO1/4 kinetics

Detailed information on the modelling process is provided in **SI Appendix**, Extended methods.

### Data representation and statistics

Detailed information is provided in **SI Appendix**, Extended methods.

## Supporting information

SI Appendix_Lavilla-Puerta

## Acknowledgements

The authors would like to thank Prof. Pierdomenico Perata for his suggestions and support during the early development of this project, and Beatriz Pasero Garcia for her help and insight on the promoter analysis. This work was partially supported by the UKRI Biotechnology and Biological Science Research Council Pioneer Award, grant CBR00770, to M.L.-P. and F.L.

