## Supplementary material for "A synthetic ERFVII-dependent circuit in yeast sheds light on the regulation of early hypoxic responses of plants": SI Appendix_Lavilla-Puerta

**This PDF file includes:**

Supporting text

Figures S1 to S10

Tables S1 to S11

SI References

### Supporting information Text

### Extended methods

Plant model description

Gene expression analysis from Col-0 seedlings growing on vertical plates subjected to 2 h hypoxia in **Fig. 1** was fitted to a logistic equation with the following formula:

$$y(t) = \frac{A}{1 + e^{-k(t-t_{half})}}$$

where $y(t)$ is the relative mRNA expression at time $t$, $A$ is the upper asymptote (maximum level of $y(t)$), $k$ is a constant related to the steepness of the curve, and $t_{half}$ is the time at which $y(t)$ is $A/2$ and is also the point where the curve grows most rapidly. All values for $A$, $k$ and $t_{half}$ were fitted to each of the nine HRGs displayed in **Fig. 1** using MATLAB's `lsqcurvefit`, and are shown in **SI Appendix, Table S1**.

To quantify the speed of HRG response, we defined response time ($RT$) as $t_{90\%}$ - $t_{10\%}$, representing the time interval required for expression to rise from 10% ($t_{10\%}$) to 90% ($t_{90\%}$) of the maximum. We then applied a linear mixed-effects model using MATLAB's `fitlme` function, to test whether the density of HRPE elements within the promoters were associated with the $RT$, while also accounting for scaled promoter length (hereafter denoted as $PromoterLength$ ):

$$RT_{ij} = \beta_0 + \beta_1 \cdot HRPEDensity_{ij} + \beta_2 \cdot PromoterLength_{ij} + u_j + \varepsilon_{ij}$$

Here $\beta_0$ is the intercept, $\beta_1$ and $\beta_2$ are fixed effects of HRPE density and scaled promoter length, $u_j$ is the random effect capturing gene-specific variation, and $\varepsilon_{ij}$ is the residual error. Because promoter lengths vary substantially among HRGs, we standardized them using z-score normalization:

$$PromoterLength_{ij} = \frac{P_{ij} - \mu}{\sigma}$$

where $P_{ij}$ is the promoter length, $\mu$ is the mean of the promoter lengths across all HRGs, and $\sigma$ is the corresponding standard deviation of the promoter lengths. This scaling places both predictor variables (HRPE density and promoter length) on comparable scales, improving interpretability and numerical stability of the regression estimates.

Yeast transformation

Yeast was transformed according to (3), with minor modifications. Prior to transformation, wild-type cells were grown at 30°C, 150 rpm on YPDA containing 20 g $L^{-1}$ peptone, 10 g $L^{-1}$ yeast extract, 20 g $L^{-1}$ of glucose (Duchefa) and 20 mg $L^{-1}$ adenine hemisulfate (Sigma-Aldrich), supplemented with 20 g $L^{-1}$ agar (Duchefa). Single colonies were grown to saturation (overnight) in 5 ml of liquid YPD, at 30°C under continuous shaking at 150 rpm, then diluted 1:100 in 50 mL liquid YPD and grown for 6 additional hours. Cells were then harvested by centrifugation at 5000 g, 5 min and washed three times with sterile water. For each transformation, 100 µL of cell suspension were centrifuged and resuspended in a buffer containing 36% PEG 3350, 100 mM lithium acetate and 100 µg single-strand DNA, plus 1 µg of plasmid DNA. Integrative plasmids were first linearized by restriction for 2 h at 37°C, with Anza endonucleases (Thermo-Fisher Scientific): pAG305-, pAG303- and pAG304-based plasmids were linearized with BstXI (cat. IVGN0666), pAG306-based plasmids with ApaI (cat. IVGN0326). Cells were incubated at 42°C for 60 min. Transformed cells were collected by centrifugation and plated on the appropriate solid SD medium (20 g $L^{-1}$ agar) for selection. Positive transformants were transferred to liquid SD, prior to further transformations or experiments. All yeast strains used in this work are listed in **SI Appendix, Table S5**.

Cloning of constructs for yeast and plant expression

Some of the Gateway-compatible destination vectors listed were purposely generated for this study, by modification of the existing destination plasmids for yeast expression. In detail, pAG304NLUC was generated from pAG304GPD-ccdb (Addgene, #14135) by restriction and ligation of a synthetic ccdb-NLUC string (GeneArt, Thermo-Fisher Scientific, sequence in **SI Appendix, Table S6**) flanked by SacI and XhoI restriction sites. pAG304GPD-ccdb and ccdb-NLUC were cut with SacI and XhoI (cat. IVGN0208 and IVGN0086, Anza, Thermo-Fisher Scientific) and joined using the Anza T4 DNA ligase (cat. IVGN2104, Thermo-Fisher Scientific). The destination plasmids $pAG303HRPE_{ADH}$ and $pAG305HRPE_{ADH}$ were obtained from pAG303GPD and pAG305GPD backbones (Addgene, #14135 and #14138, respectively), produced upon removal of P*GPD* promoter sequence following a digestion with SacI and XbaI (IVGN0126, Anza, Thermo-Fisher Scientific). The promoter was replaced by ligation with an $HRPE_{ADH}$ sequence, amplified from $HRPE_{ADH}$_pE (**SI Appendix, Table S8**) with compatible SacI and XbaI extremities incorporated in the PCR primers (**SI Appendix, Table S7**). The destination vector pBGWNLUC7 was, instead, obtained from pBGWL7 (4) and used to generate the binary construct $HRPE_{ADH}$-NLUC-pBGWNLUC7 for stable plant transformation. A DNA string containing the SacI-attR1-ccdB-attR2-NanoLuc-HindIII sequence was synthesized (GeneArt, Thermo-Fisher Scientific; **SI Appendix, Table S6**) and recombined into the pBGWL7 upon restriction with SacI and HindIII (IVGN0168, Invitrogen) and ligation with Anza T4 DNA ligase. Recombinant plasmids were propagated *E. coli* One Shot *ccdB* Survival 2 $T1^R$ Competent Cells (Thermo-Fisher Scientific), before use in LR reactions.

PCO4-SV40 was cloned from 7-day-old Arabidopsis seedling cDNA with two subsequent PCR reactions, using two overlapping primers (**SI Appendix, Table S7**) to incorporate a C-terminal SV40 domain (PKKKRKV) into the PCO4 coding sequence. To generate the RAP2.12-derived chimeric transcription factors, the full RAP2.12 coding sequence was amplified from the same cDNA template as above and fused to different 3' fragments by overlapping PCR. Specifically, RAP2.12 coding sequence was amplified from Arabidopsis cDNA from seedlings and fused to a 6xTAL-4xVP16 fragment amplified from the plasmid dCas9-TV (5) to obtain RAP2.12-6TVP. The fusion sequence was created by overlapping PCR of fragments amplified with a combination of RAP2.12_Fw and RAP2.12-6TVP overlapping primers from **SI Appendix, Table S7**. Similarly, a Gal4 AD was amplified from the plasmid pDEST22 (Thermo-Fisher Scientific) and fused to RAP2.12 to obtain the RAP2.12-GAL4AD insert. In the case of SYRAP, finally, a chimeric GAL4Ste sequence ($GAL4_{1-147}$-$Ste12_{301-335}$-$GAL4_{148-196}$) was fused, using SYRAP overlapping primers. The GAL4Ste template (6) was provided by a synthetic NLUC-UBQ-$RAP2.12_{2-50}$-GAL4STE string (**SI Appendix, Table S5**), purchased from GeneArt (Thermo-Fisher Scientific). The ubiquitin fusion factor UbSYRAP was further amplified from the entry plasmid UbSYRAP_201 SYRAP. A UBQ fragment was amplified from the same string DNA template, with SY_UBQ_Fw and OL_SY_Rv primers, and combined with a SYRAP fragment amplified with OL_SY_Fw and G4Ste_Rv (**SI Appendix, Table S5 and S7**) UbSYHRE was generated by overlapping PCR between the synthetic templates UBQ-HRE2 and GAL4Ste (**SI Appendix, Table S6**).

All above mentioned constructs were made BP-compatible thanks to the presence of BP adaptors in their flanking sequences. The inserts were subsequently amplified with primers BP_Ad_Fw and BP_Ad_Rv (**SI Appendix, Table S7**), to restore full attB sites, and recombined into pDONR201 using the Gateway™ BP Clonase™ II Enzyme mix (Thermo-Fisher Scientific).

$UAS_{GAL}$ activation experiments in MaV203 strains

Overnight cultures were diluted to $OD_{600}$ = 0.1 and grown for 6 h. Growth rates were calculated from optical density values of the microcultures, measured using the Multiskan Go 1510 Sky plate reader (Thermo-Fisher Scientific). To overcome the low saturation limit of optical detectors in microplate readers, the $OD_{600}$ values were corrected by means of an empirical formula (7), where (corrected $OD_{600}$) = $-8.7202(OD_{600})^2 + 11.363(OD_{600})$. Cells were then diluted to $OD_{600}$ = 0.1 and re-incubated at 30°C, to follow their growth at 2 h intervals. Growth rate (k) was calculated between 4 and 6 h after dilution, when positive controls showed a clear exponential growth phase (**SI Appendix, Fig. S4**), according to the following formula (8):

$$k = \frac{log_{10}\left(\frac{corrected\ OD_{600,6h}}{corrected\ OD_{600,4h}}\right)}{0.301 \cdot 2}$$

Β-galactosidase (LacZ) activity from the P*GAL1-LacZ* reporter was measured by adapting to yeast a colorimetric assay initially developed for animal tissues (9). Approximately $2x10^7$ cells were collected by centrifugation, washed twice with sterile water, once with PBS (137 mM NaCl, 2.6 mM KCl, 10 mM $Na_2HPO_4$, 1.8 mM $KH_2PO_4$; pH=7,4) and resuspended in a buffer composed of 0.25 M Tris-HCl, 2.5 M EDTA, 0.2 M lithium acetate, 1% v/v Triton X-100 and 4.2 µl $mL^{-1}$ antifoam (Merck) (pH=7.4). Cell suspensions were flash frozen in $N_2$ and then homogenized with a Bead Mill 4 Homogenizer (Fisher Scientific), using two cycles of 120 sec at maximum speed. For the colorimetric assay, 100 µl of the lysate were mixed with 900 µl of CPRG solution containing 25 mM MOPS, 100 mM NaCl, 10 mM $MgCl_2$ (pH=7.5) and 84 mg $l^{-1}$ of red-β-d-galactospyranoside (CPRG) substrate (Merck). Samples were incubated at 37°C and monitored until a color shift from orange to red became visible (8 or 24 h), indicative of substrate hydrolysis. CPRG absorbance at 575 nm was measured in a spectrophotometer and LacZ activity was calculated according to (9).

To prepare spots, $10^5$ cells were collected, washed with 1 ml sterile water four times and diluted on four 1:10 serial dilutions. Suspensions were spotted in 10 µl volume on -leu-trp SD solid media supplemented with 10 mM 3-amino-1,2,4-triazole (3-AT, Merck), found to be the optimal concentration to overcome the leaky HIS3 activity in the negative controls (EV/RAP2.12), while allowing growth of the positive controls expressing GAL4Ste. Plates were incubated for 3 days at 30°C either in normoxic or hypoxic conditions and scanned.

Immunoblotting

Total protein samples were extracted from $2\text{-}3x10^8$ yeast cells in late exponential cultures (10). Arabidopsis seedling extracts were prepared according to (11). A positive control for GAL4 immunoblottings was provided by a recombinant protein, produced *in vitro* via the TnT Coupled Wheat Germ Extract Systems (Promega). To this end, the NLUC-UBQ-$RAP2.12_{2\text{-}50}$-GAL4Ste string (**SI**

**Appendix, Table S6**) was LR cloned into the pF3A WG (BYDV) Flexi Vector (Promega) and *in vitro* transcription and translation was performed, following the manufacturer's recommendations.

Proteins were separated via SDS-PAGE on 10% polyacrylamide gels (NuPage Bis-Tris Gels, Thermo-Fisher Scientific) and transferred to poly-vinylidene difluoride membranes (Bio-Rad), as described before (2). Yeast blots were hybridized with primary monoclonal GAL4-DBD (sc-577) or α-tubulin antibodies (sc-53030, Santa Cruz Biotechnologies), used at 1:1000 and 1:5000 respectively. Plant blots were hybridized with the primary monoclonal antibodies anti-HA (#26183, Thermo Fisher Scientific, 1:1000) and anti-actin11 (AS13 2640, Agrisera 1:2000), or the polyclonal antibodies anti-ADH and anti-PDC (AS10 685 and AS10 691 respectively, Agrisera 1:1000). HRP-conjugated secondary antibodies were used for the immunodetection. A polyclonal anti-mouse IgG (BioVision, cat. no. 6402-05, 1:20000) was used against GAL4-DBD and HA; anti-Rat IgG (AS10 1187, Agrisera, 1:10000) against α-tubulin, and anti-Rabbit IgG (AS09 602, Agrisera, 1:10000) for actin, ADH and PDC. Bands were visualized with a Clarity chemiluminescent substrate (1705060, Bio-Rad), using a ChemiDoc MP Imaging System (Bio-Rad). Protein loading on the gels was evaluated by membrane staining with 0.1% amido black 10-B (2). Band intensity quantification was performed with the ImageLab® software (Bio-Rad).

Yeast model description and assumptions

To build the four Ordinary Differential Equation (ODE)-based models illustrated in **Fig. 6A**, we incorporated both Michaelis-Menten Kinetics and Mass Action laws to describe the gene expression and biochemical reaction network processes respectively (**SI Appendix, Tables S2** and **S3**).

The basic model (GG) represents a single copy of a constitutively expressed RAP2.12 and PCO. Here, RAP2.12 (UbSYRAP) protein is synthesized at a rate $k_1$ and degraded at a rate $k_2$. Additionally, it undergoes degradation by the cysteine N-degron pathway, following Michaelis-Menten kinetic laws, at rates represented by $k_3$, $K_{m1}$, and $K_{m2}$. The activation of HRPE promoter, which is targeted by RAP2.12 depending on the binding rates of RAP2.12 to the HRPE promoter for NLUC/FLUC ($HRPE_1$), produces NLUC/FLUC activity following Michaelis-Menten kinetics at a rate represented by $k_4$ and $Km_3$ and NLUC/FLUC undergoes degradation at a rate of $k_5$. In the models with additional feedback loops involving PCO1 (GP), RAP2.12 (UG), and both (UP), the HRPEs are targeted by RAP2.12 depending on the binding rates of RAP2.12 to the HRPE promoter for the PCO1 ($HRPE_2$) and RAP2.12 ($HRPE_3$) feedback loop, respectively. Subsequently they are activated following Michaelis-Menten kinetics at rates represented by $k_6$ and $k_9$, respectively and $Km_3$. PCO1 has unique Michaelis-Menten kinetic parameters in N-degron pathway, different from PCO4, which are $k_8$, $K_{m4}$ and $K_{m5}$. $k_4$, $k_6$ and $k_9$ are aggregated parameters overseeing all processes involved in protein synthesis rate.

To simplify our model, each reaction is assumed to take place in a spatially homogeneous environment, without considering compartmentalization or specific microenvironments within the cell and that each reaction species, including RAP2.12, PCO1/4, oxygen, and NLUC, is evenly distributed. To concentrate on the oxidation process and simplify the model, we only modelled RAP2.12 degradation after its oxidation by PCO1/4, assuming arginylation, ubiquitination, and degradation by ATE, PRT6, and the 26S proteasome happen immediately afterwards. Additionally, we assumed that the concentration of PCO4 has already reached a steady state during normoxia and remains constant even after applying the 1% hypoxia treatment. Thus, PCO4 can be considered as a parameter without introducing its own expression and degradation rates which simplifies and adds more flexibility to our model.

Non-linear fitting and simulation

We used MATLAB's 'Talwar' and 'Welsch' weight functions within the `nlinfit` function for robust fitting to re-fit the catalytic rate constants ($k_8/k_3$) for PCO1/4-mediated oxidation of RAP2.12, the Michaelis-Menten constants ($K_{m4}/K_{m1}$) for $O_2$ as a substrate of PCO1/4, and the Michaelis-Menten constants ($K_{m5}/K_{m1}$) for RAP2.12 as a substrate of PCO1/4, respectively. This three-dimensional fitting, which is depicted in **SI Appendix, Fig. S9A**, was performed based on published RAP2.12 and $O_2$ kinetic data (1). The initial guesses for RAP2.12 protein synthesis rate ($k_1$) and degradation rate ($k_2$), and PCO4 concentration in yeast were obtained by simulation visualization using MATLAB's ode45 solver, and subsequently the parameters were fitted using the `lsqcurvefit` function, initialised by those guesses, based on the published yeast data with proper rescaling (2). (**Fig. S9B**). Additionally, we applied the same rate for pGDP-regulated RAP2.12 production and degradation in all four different models. The remaining parameters in the models were fitted using the same method, based on the data shown in **Fig. 5C**. All parameters are listed in **SI Appendix, Table S4**.

To simulate hypoxic responses, the initial concentrations of RAP2.12, PCO1, and mRNA as inputs were obtained from steady-state simulations conducted under normoxic conditions (21% $O_2$) for six hours (**SI Appendix, Tables S2 and S3**). Subsequently, dynamic simulations of hypoxic responses (NLUC/FLUC) and adjusted RAP2.12 (log10(RAP2.12)/2) were conducted under 1% $O_2$ conditions for four hours.

Data representation and statistics

Box plots represent median (line) and interquartile range (IQR), whiskers span from minimum to maximum values. Graphpad Prism v. 8.0.1 was adopted to depict data and to evaluate statistical significance, which passed a Shapiro-Wilk normality test ($p<0.05$) prior to the analysis. Pairwise comparisons through Student's t-test (n=4-6) were applied to data in Figure 1C, Figure 2B, Figure 3B and C, Figure 4 and Figure 5C, as well as in SI Appendix, Figures S1 and C, S3A, S4B and C, S5, S6B, D and E, and S8. For the analysis on figures S6D and S8, separate analysis tables were generated as **SI Appendix, Table S10** and **Table S11**. One-way Anova was used in Figures 2D, 3A and E, 5 E and F and **SI Appendix**, S6E (in these last three, treatment-independent analysis). In both kinds of procedure, the multiple comparisons were corrected with a two-stage linear step-up procedure of Benjamini, Krieger and Yekutieli, with Q=1%. Statistical significance was depicted as asterisks (****, $p<0.0001$; ***, $0.0001\leq p<0.001$; **, $0.001\leq p<0.01$; *, $0.01\leq p<0.05$) or letters (for $p<0.05$, unless stated otherwise). Two-way Anova was, instead, adopted in and Figures S3B, S6A and F, where the analysis was followed by a Tukey post-hoc test, with letters depicting significantly different groups for $p<0.05$.

Grey shadings in figures indicate statistical difference, according to multiple Student's t-test, between a given time point in hypoxia and the normoxic control ($t_0$). The first group to be significantly different from normoxia ($t_0$), regardless of further changes, was shaded in Figure 1A and 5B, and **SI Appendix Figures** S1A and C, S4A and C, S5A and C and S6B. Instead, where different shades of grey appeared, they depicted different significance levels corresponding to the accompanying asterisks (****, $p<0.0001$; ***, $0.0001\leq p<0.001$; **, $0.001\leq p<0.01$; *, $0.01\leq p<0.05$) in Figure 4C-F, 5C and **SI Appendix**, Figure S6B.

### Supporting Figures

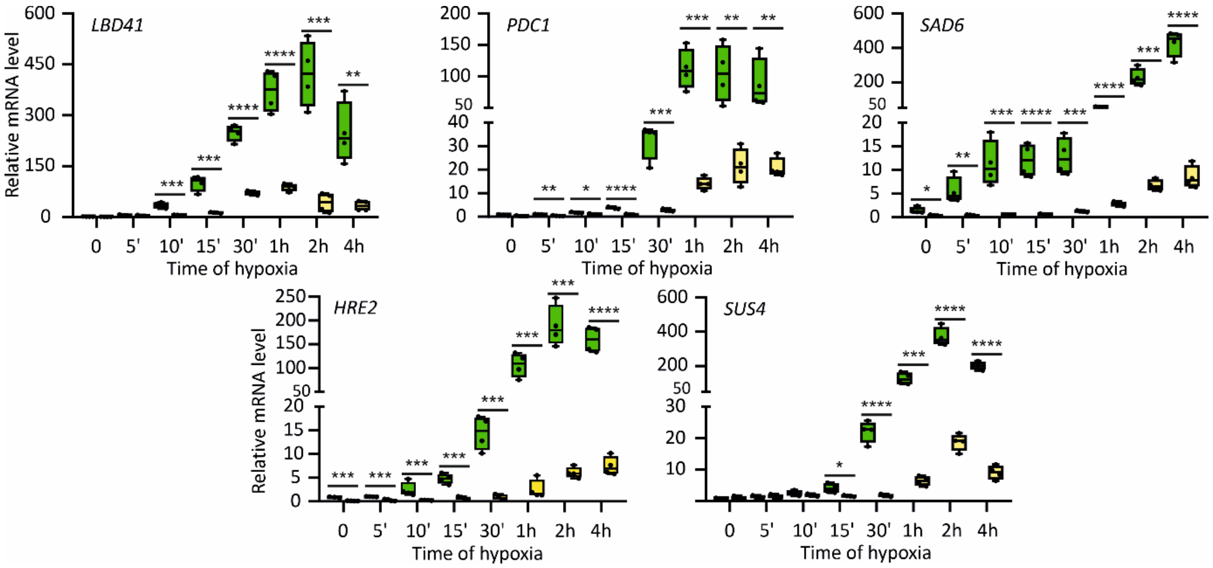


**Figure S1. Profiling of hypoxia-responsive gene (HRG) expression in Arabidopsis *erfvii* mutants**. *Lateral organ Binding Domain 41* (*LBD41*), *Pyruvate Decarboxylase 1* (*PDC1*), *Steraroyl-Acyl Carrier Protein 6* (*SAD6*), *Hypoxia Responsive ERF 2* (*HRE2*) and *Sucrose Synthase 4* (*SUS4*) transcript abundance (normalized to *UBQ10* and relative to an aerobic Col-0 sample) in *erfVII* and Col-0 7-day-old seedlings, exposed to dark hypoxia (1% $O_2$ v/v) at 23°C, for 4 hours. Asterisks indicate significant differences after pairwise comparisons between genotypes, forn=4 (****, p<0.0001; ***, 0.0001≤p<0.001; **, 0.001≤p<0.01; *, 0.01≤p<0.05).

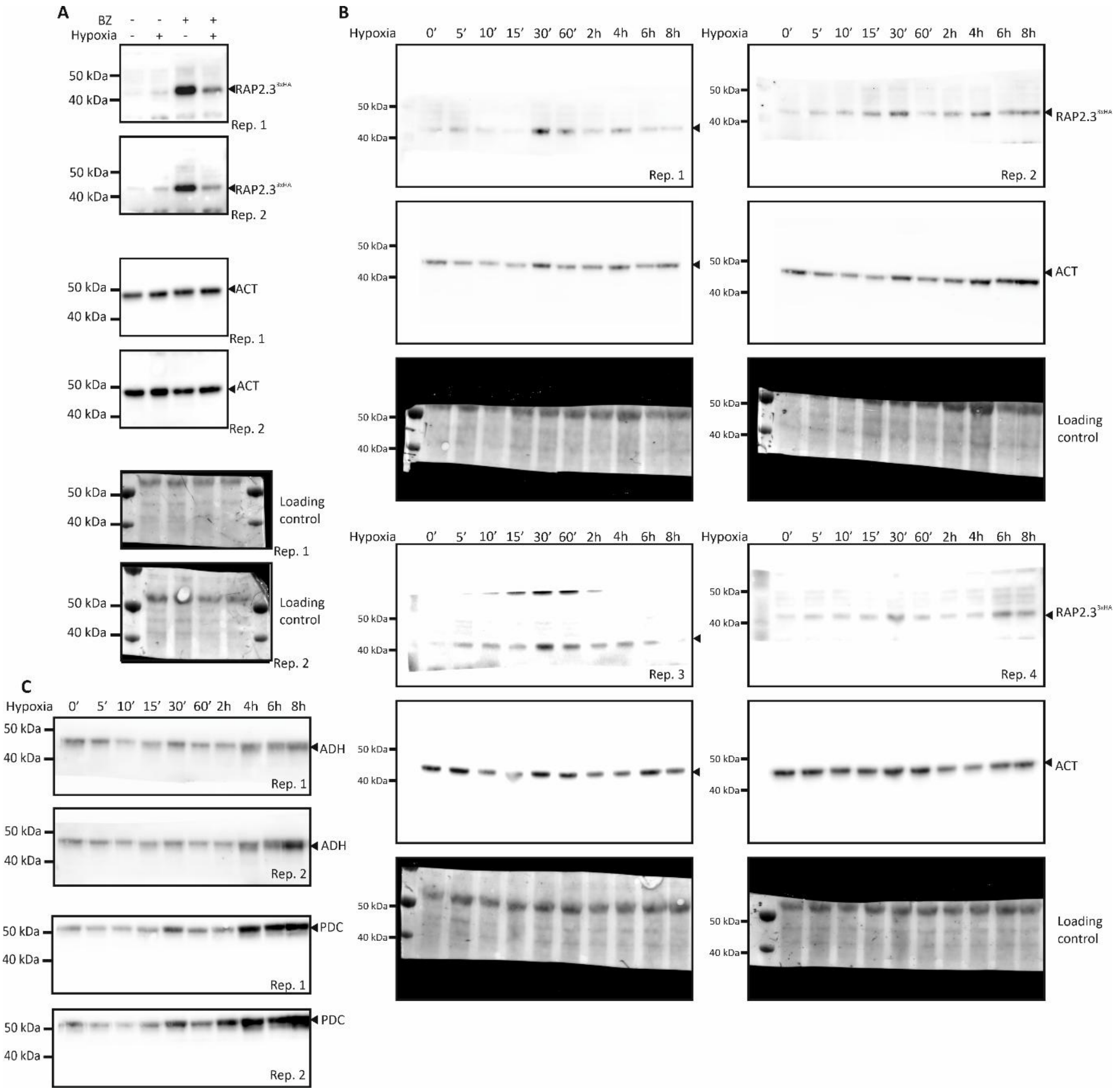


**Figure S2. Uncropped blots from Arabidopsis protein samples. (A)** Supporting blots for Figure 2B. Seedlings were grown in liquid medium for 7 days and then treated with 100 µM bortezomib (BZ) or 1% (v/v) DMSO. **(B)** Supporting blots for Figure 2D. Detection of HA-tagged RAP2.3, actin (ACT), and loading control (n=4) replicates. **(C)** Blots showing alcohol dehydrogenase (ADH) and pyruvate decarboxylase (PDC) in replicates 1 and 2 from panel (B). All proteins were detected from total protein extracts of 7-day-old Arabidopsis seedlings containing a *35S:RAP2.3$^{3xHA}$* transgene, grown in vertical plates and moved from normoxia ($t_0$) to hypoxia (1% $O_2$ v/v) for the indicated duration.

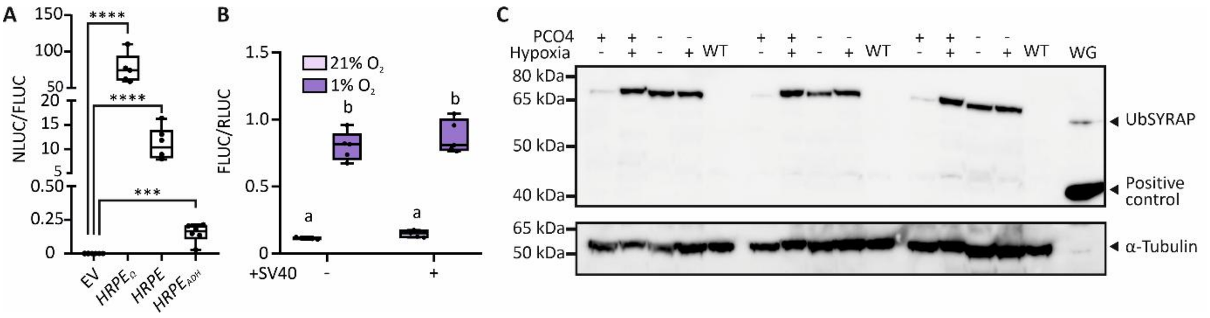

**Figure S3. Characterization of the components of the yeast transcriptional circuit**. **(A)** Basal NLUC activity of *HRPE-NLUC, $HRPE_{ADH}$-NLUC* and *$HRPE_{\Omega}$-NLUC* promoter modules, in comparison with an empty vector (EV) (n=5-6). Output signals are expressed as ratios between NLUC activity and FLUC activity from a reference *GPD (glyceraldehyde 3-phosphate dehydrogenase) promoter: FLUC* integrative construct. Output signals are expressed as activity ratios with FLUC encoded by a reference P*GPD:FLUC* integrative construct. **(B)** Activity of the SV40-tagged PCO4 version used in this study, expressed by the integrative plasmid pAG305 (12), on the model substrate DLOR (2). DLOR output is displayed as FLUC/RLUC ratio (n=5). For reference, the untagged PCO4 version used in (2), expressed by the episomal plasmid pAG415 (12) was included in the experiment. **(C)** Uncropped blot of yeast cells expressing UbSYRAP and PCO4 or GUS (-PCO4) treated with hypoxia. A triplicate experiment is shown. UbSYRAP was revealed with an α-GAL4BD antibody. α-Tubulin was used as housekeeping protein on the same stripped and hybridized membrane. WT, untransformed W303 cells. WG, wheat germ extract expressing a recombinant NLUC-UBQ-RAP2.$12_{2\text{-}50}$-GAL4STE12 protein, as positive control. (**SI Appendix, Table S6**). Asterisks in (A) represent statistically different groups, as tested via multiple Student's t-tests (n=6; ****, p<0.0001; ***, 0.0001≤p<0.001). Letters in (B) represent statistically different groups, as tested by two-way Anova, for n=5 and p<0.05.

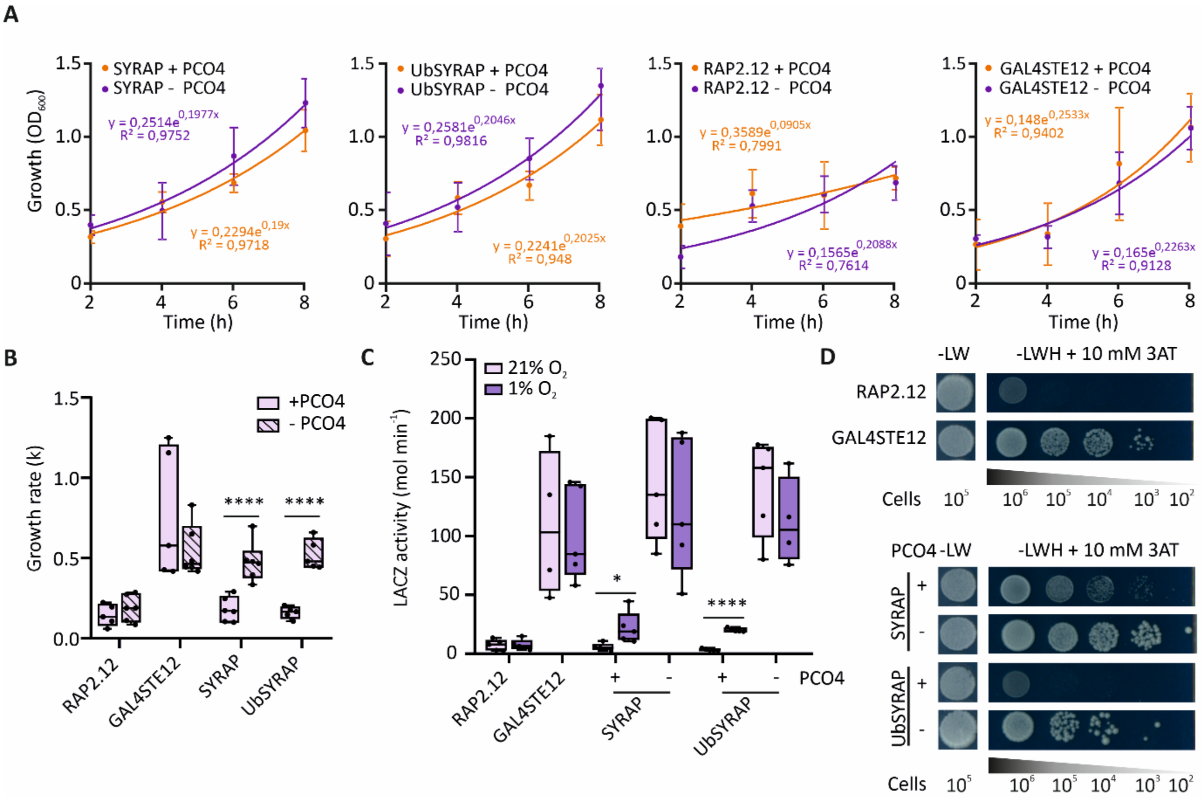


**Figure S4. Orthogonal Cys-NDP regulation in MaV203 strain yeast**. **(A)** Growth curves ($OD_{600}$) of aerated cultures expressing SYRAP or UbSYRAP, along with either PCO4 or a β-glucoronidase (GUS) protein (-PCO4). GAL4STE or RAP2.12 expressing colonies were also tested as positive or negative controls, respectively. GAL4STE can autonomously bind and activate $UAS_{GAL}$, while RAP2.12 lacks a GAL4-BD domain. Exponential regressions were fitted to the data, displayed as mean ± SD (n=5). **(B)** Growth rate constants (k) calculated between $t_6$ and $t_4$ from (A). **(C)** LACZ activity in liquid cultures after 6 h hypoxic treatment. LACZ activity was measured through colorimetric CPRG assay. **(D)** Aerobic growth of spotted colonies on –leu-his-trp (-LWH) solid media, supplemented with 10 mM 3-AT. Ten-fold serial dilutions were made to the approximate cell number indicated below. Asterisks represent statistically different groups, as tested via paired Student's t-tests, for +PCO4 vs. -PCO4 (B) and air vs. hypoxia (C); (n=4-6; ****, p<0.0001; *, p<0.05)

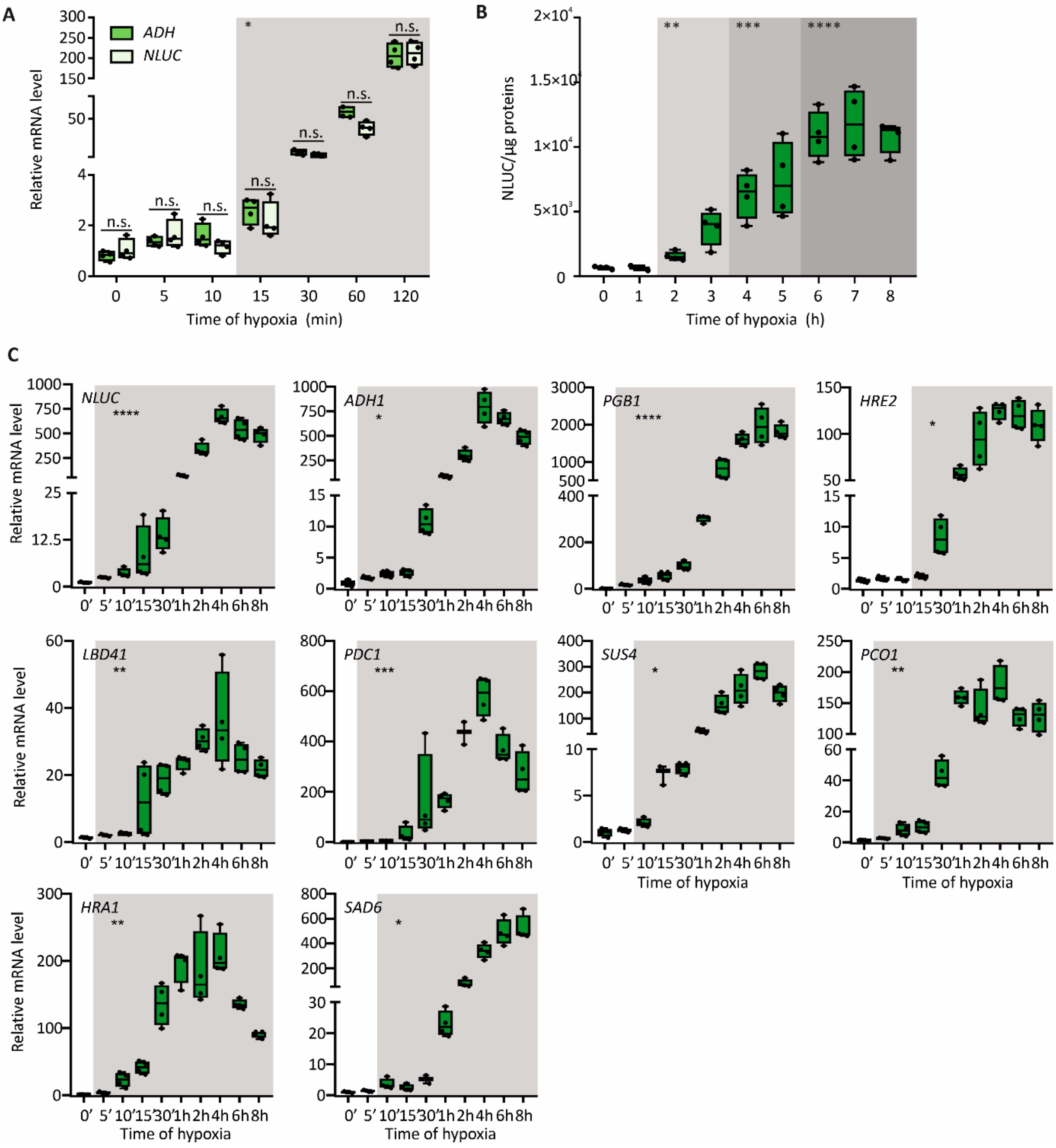


**Figure S5. Characterization of *NLUC* expression and long-term activity in the HRPE reporter line of Arabidopsis. (A)** Relative expression to a normoxic sample of *ADH1* and *NLUC,* normalized to *UBQ10*. Paired Student's t-tests for each time-point were performed (n=4; n.s., p≥0.05). **(B)** Long-term NLUC activity, normalized on total protein content, in seedlings exposed to a time course of hypoxia. **(C)** Extended profiling of HRG expression in the same plants as (B). All experiments were carried out with 7-day-old *HRPE$_{ADH}$-NLUC* seedlings grown on vertical plates. Asterisks and shading indicate statistically significant differences after multiple Student's t-tests between each time-point and t0 (n=3-4; ****, p<0.0001; ***, 0.0001≤p<0.001; **, 0.001≤p<0.01; *, p<0.05) ; in (C), only the first statistically different group was highlighted.

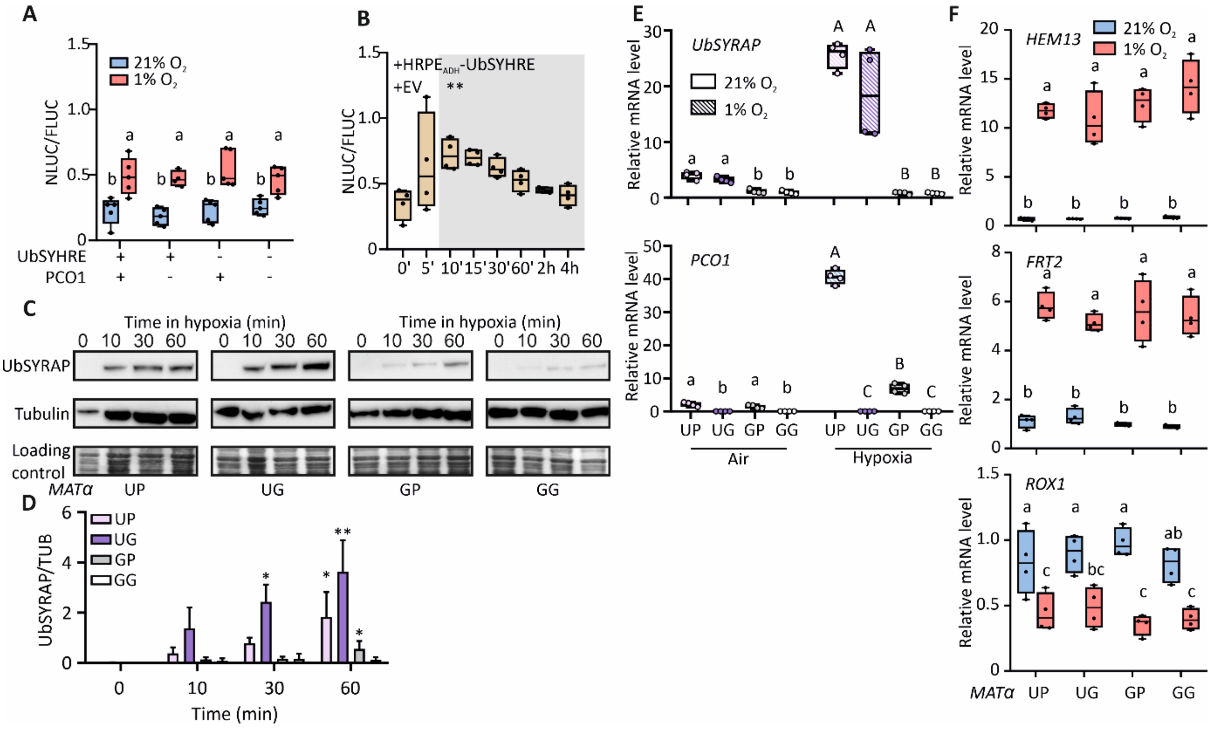


**Figure S6. Characterization of the hypoxic feedback loop in yeast. (A)** NLUC activity (normalized to P*GPD:FLUC*) in MynOx$^{MATa}$ cells mated with *MATα* cells containing combinations of $HRPE_{ADH}$-UbSYHRE, $HRPE_{ADH}$-PCO1 or $HRPE_{ADH}$-GUS (“-“). Cultures were incubated for 6 h in hypoxia or normoxia. **(B)** NLUC activity in diploid MynOx$^{MATa}$/$HRPE_{ADH}$-UbSYHRE$^{MATα}$ cells, growing on solid phase, challenged with up to 4 h hypoxia. **(C)** UbSYRAP protein abundance and **(D)** quantification (normalized to α-tubulin) (n=3-4) in the four diploid strains during a “hypoxic chase”. Cultures were treated with 1% $O_2$ in pre-equilibrated media for up to 60 min. Replicate blots are provided in **SI Appendix, Figure S7.** Asterisks indicate significant differences after Studdent’s t-test and pairwise comparisons between a given genotype and GG for the same time-point (n=3-4). Full statistical multiple analysis is provided in **Table S10**. **(E)** *UbSYRAP* and *PCO1* mRNA levels in diploid colonies before (21% $O_2$) or after 1 h hypoxia at 1% $O_2$. Different letters indicate statistically significant differences after One-way Anova in normoxic (lowercase) or hypoxic (uppercase) samples (n=4, $p<0.05$) with Benjamini, Krieger and Yekutieli correction. **(F)** Expression of yeast hypoxia marker genes in diploid colonies treated with 1% $O_2$ for 4 hours. In (A) and (F), letters indicate statistically significant differences after Two-way Anova (n=4-5, $p<0.05$) followed by Tukey post-hoc test.

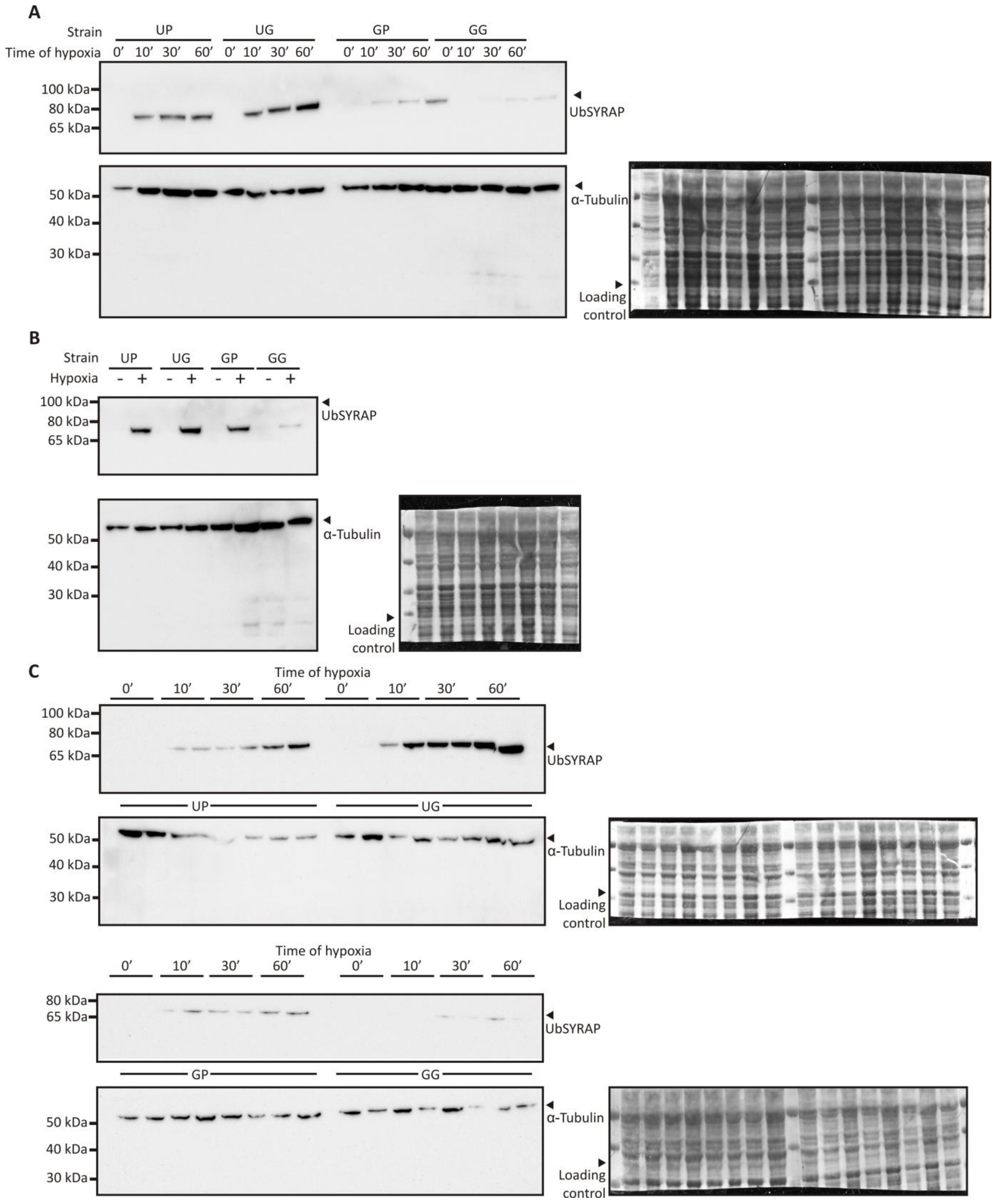


**Figure S7. Supporting blots for Figure 4 and SI Appendix, Figure S6.** UbSYRAP and α-tubulin protein abundance in MynOx$^{MAT\alpha}$ cells mated with *MATα* strains containing HRPE$_{ADH}$-UbSYRAP (U), HRPE$_{ADH}$-PCO1 (P) or HRPE$_{ADH}$-GUS (G). Up to 3 blots **(A, B, C)** with different samples were used for quantification. Amido black stained membranes are shown besides the corresponding blots to reveal total protein loading.

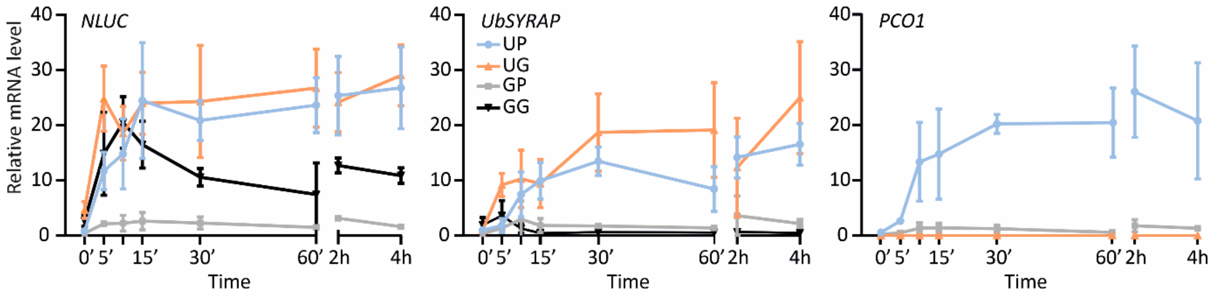


**Figure S8. Solid-phase mRNA dynamics in yeast**. Profiling of *NLUC, UbSYRAP* and *PCO1* expression in diploid cells grown in thin spots (pre-grown for 28 h on solid phase) over a 4 hour-long time course of hypoxia (1% $O_2$). Supporting statistics in **Table S11**.

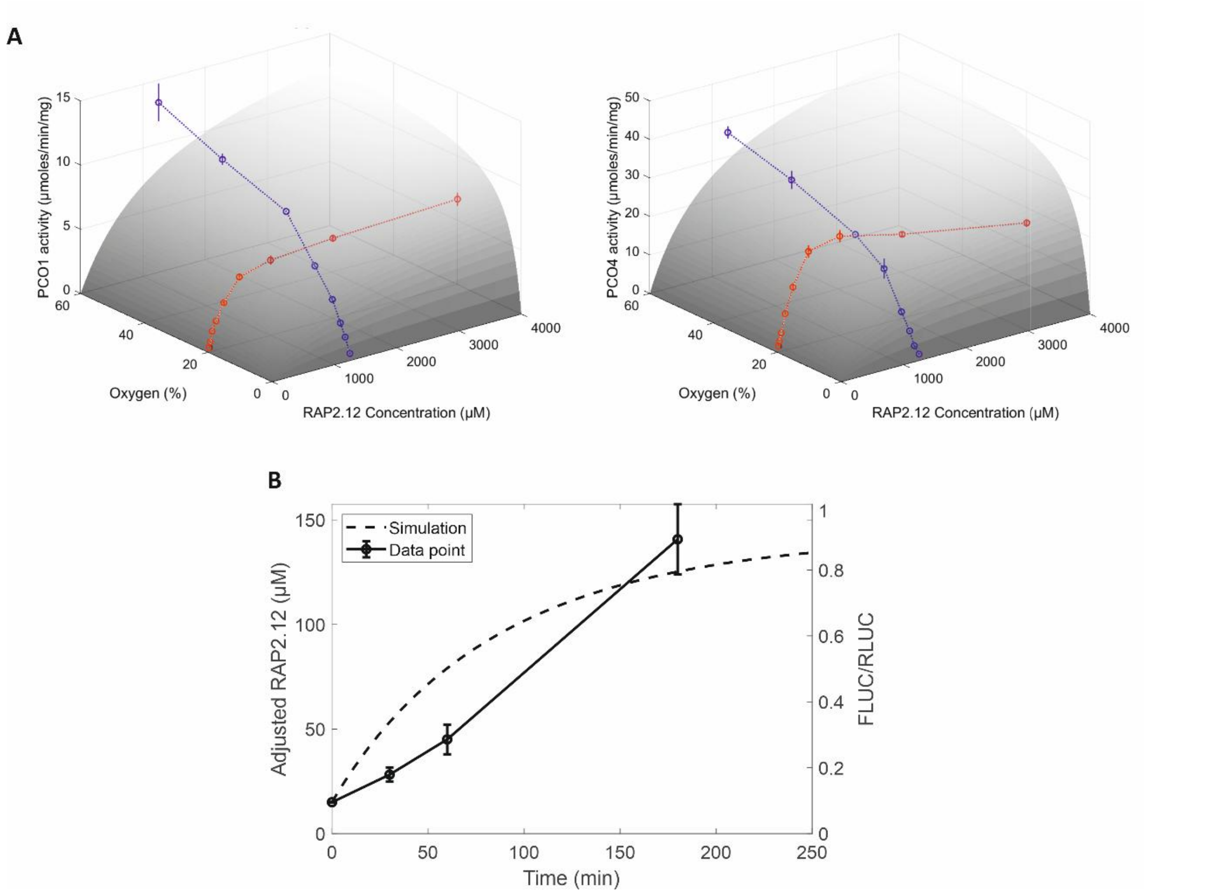


**Figure S9. Kinetic parameters fitting and visualization. (A)** 3-D data visualization and fitting for RAP2.12 and PCO1/4 kinetics. Kinetic data from (1), with blue and red lines indicating PCO activity at different oxygen and RAP2.12 concentrations, respectively. **(B)** Kinetic parameters fitting for RAP2.12 synthesis rate by the *GPD* promoter and degradation rate in yeast. RAP2.12 (UbSyRAP) concentration was rescaled based on published DLOR data from (2).

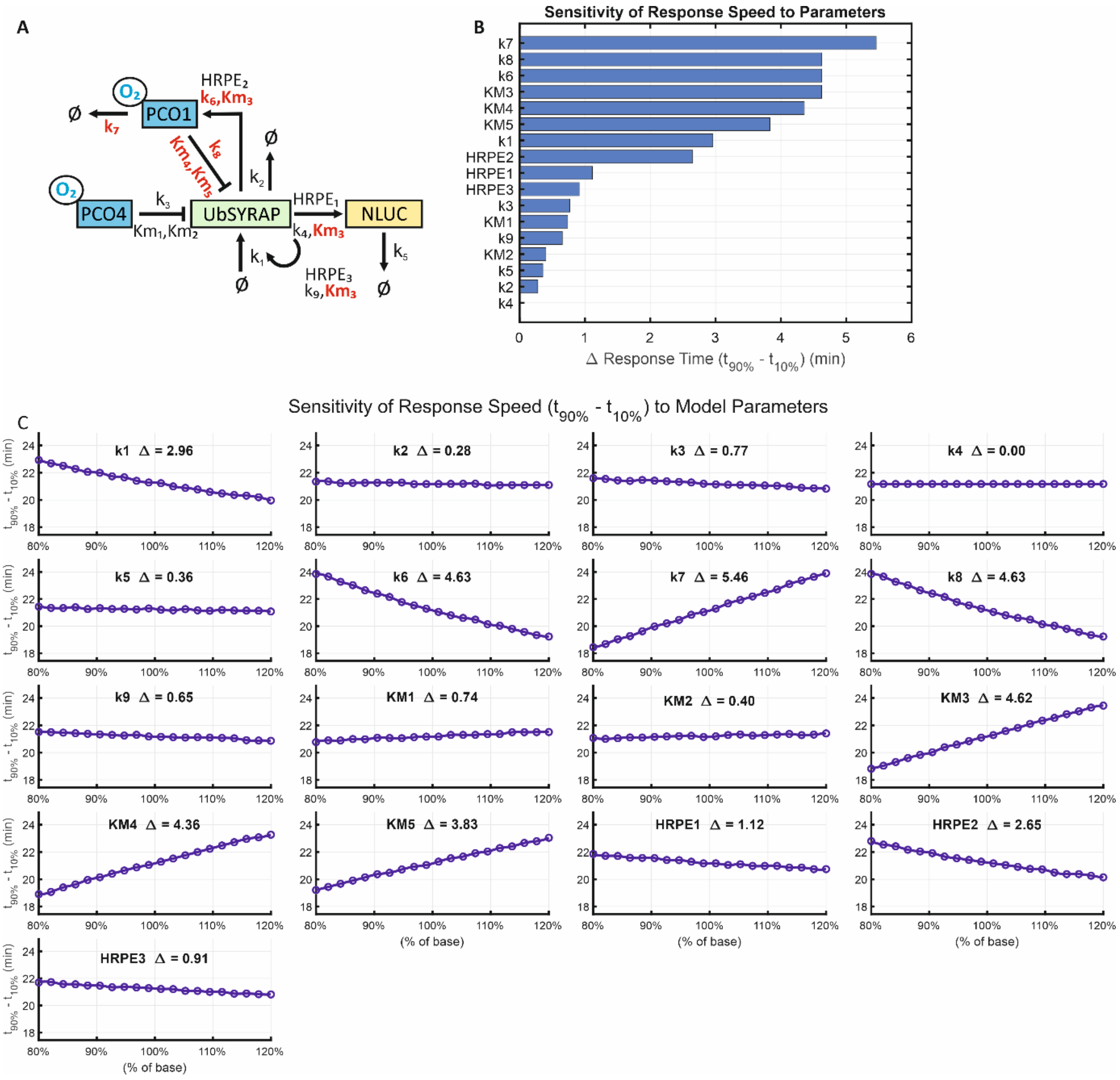


**Figure S10. Response time sensitivity analysis to the model parameters. (A)** UP model, highlighting in red the top 6 parameters affecting the model's Response time (RT; $t_{90\%}$-$t_{10\%}$). **(B)** Ranking of parameters affecting RT. **(C)** Single parameter sensitivity analysis. Parameters were iterated from 80% to 120% their fitted value (% of base).

### Supporting Tables

**Table S1. Parameters used for HRG expression equations**. Parameters defining the logistic equation fitted into gene expression analysis of Fig. 1A (2 h hypoxia, 1% v/v $O_2$ in Col-0 seedlings growing in vertical plates).

| Gene | Max. expression ($A$) | Rate constant ($k$) ($min^{-1}$) | Half-max time ($t_{half}$) (min) |
|---|---|---|---|
| *ADH1* | 326.32 | 0.064 | 74.87 |
| *PGB1* | 83.27 | 0.046 | 74.46 |
| *HRE2* | 219.14 | 0.076 | 51.46 |
| *SAD6* | 137.93 | 0.052 | 87.59 |
| *PCO1* | 127.09 | 0.092 | 31.94 |
| *HRA1* | 171.66 | 0.317 | 12.39 |
| *LBD41* | 229.40 | 0.302 | 11.26 |
| *PDC1* | 145.89 | 0.060 | 54.43 |
| *SUS4* | 103.73 | 0.068 | 63.71 |

**Table S2. Ordinary differential equations of the RAP2.12 and PCO1/4 kinetic model.** The reaction rates v1 - v9 are given in Table S3.

| | Left-hand Sides | Right-hand Sides | Initial Concentrations (Simulated) |
|---|---|---|---|
| GG model | d[RAP2.12]/dt | v1-v2-v3 | 9.1087 µM |
| | d[NLUC/FLUC]/dt | v4-v5 | 0.2402 |
| GP model | d[RAP2.12]/dt | v1-v2-v3-v8 | 3.9663 µM |
| | d[NLUC/FLUC]/dt | v4-v5 | 0.1800 |
| | d[PCO1]/dt | v6-v7 | 0.2450 µM |
| UG model | d[RAP2.12]/dt | v1-v2-v3+v9 | 10.2490 µM |
| | d[NLUC/FLUC]/dt | v4-v5 | 0.2744 |
| UP model | d[RAP2.12]/dt | v1-v2-v3-v8+v9 | 7.9160 µM |
| | d[NLUC/FLUC]/dt | v4-v5 | 0.2116 |
| | d[PCO1]/dt | v6-v7 | 0.0082 µM |

**Table S3. Reaction rates of the RAP2.12 and PCO1/4 kinetic model.** Null indicates degradation.

| | Reactions | Reaction Rates | Description |
|---|---|---|---|
| $v_1$ | → RAP2.12 | $k_1$ | RAP2.12 synthesis |
| $v_2$ | RAP2.12 → null | $k_2 \cdot [RAP2.12]$ | RAP2.12 degradation |
| $v_3$ | RAP2.12 + $O_2$ + PCO4 → PCO4 | $k_3 \cdot [PCO4] \cdot \frac{[O_2]}{Km_1+[O_2]} \cdot \frac{[RAP2.12]}{Km_2+[RAP2.12]}$ | RAP2.12 degradation via NDP by PCO4 |
| $v_4$ | RAP2.12 + $HRPE_1$→ NLUC/FLUC + RAP2.12 + $HRPE_1$ | $k_4 \cdot \frac{[RAP2.12] \cdot HRPE_1}{Km_3+[RAP2.12] \cdot HRPE_1}$ | NLUC/FLUC production by RAP2.12 binding to HRPE |
| $v_5$ | NLUC/FLUC → null | $k_5 \cdot [NLUC/FLUC]$ | NLUC/FLUC degradation |
| $v_6$ | RAP2.12 + $HRPE_2$ → PCO1 + RAP2.12 + $HRPE_2$ | $k_6 \cdot \frac{[RAP2.12] \cdot HRPE_2}{Km_3+[RAP2.12] \cdot HRPE_2}$ | PCO1 synthesis via feedback loop |
| $v_7$ | PCO1 → null | $k_7 \cdot [PCO1]$ | PCO1 degradation |
| $v_8$ | RAP2.12 + $O_2$ + PCO1 → PCO1 | $k_8 \cdot [PCO1] \cdot \frac{[O_2]}{Km_4+[O_2]} \cdot \frac{[RAP2.12]}{Km_5+[RAP2.12]}$ | RAP2.12 degradation via NDP by PCO1 |
| $v_9$ | RAP2.12 + $HRPE_3$ → RAP2.12 + RAP2.12 + $HRPE_3$ | $k_9 \cdot \frac{[RAP2.12] \cdot HRPE_3}{Km_3+[RAP2.12] \cdot HRPE_3}$ | RAP2.12 synthesis via feedback loop |

**Table S4. Parameter values used in the RAP2.12 and PCO1/4 kinetic model.** Concentrations and the Michaelis-Menten constants (Kms) are given in µM, except Kms for $O_2$ are in %. Rate constants (ks) are expressed in $s^{-1}$, except $k_1$, $k_6$ and $k_9$ are in µM $s^{-1}$.

| Parameters | Description | Values | References |
|---|---|---|---|
| $k_1$ | RAP2.12 protein synthesis rate | 0.0301 | Fitted (Puerta et al., 2019) |
| $k_2$ | RAP2.12 degradation rate | 0.0001 | Fitted (Puerta et al., 2019) |
| $k_3$ | Catalytic rate constant for PCO4-mediated oxidation of RAP2.12 | 26.5878 | Re-fitted (White et al., 2018) |
| $K_{m1}$ | Michaelis-Menten constant for $O_2$ as a substrate of PCO4 | 16.4217 | Re-fitted (White et al., 2018) |
| $K_{m2}$ | Michaelis-Menten constant for RAP2.12 as a substrate of PCO4 | 270.2505 | Re-fitted (White et al., 2018) |
| PCO4 | PCO4 protein concentration | 0.06-0.08 | Fitted |
| $k_4$ | NLUC protein synthesis rate constant measured by RAP2.12 activating $HRPE_1$. | 0.006-0.0088 | Fitted |
| $HRPE_{1GG}$ | The binding rate of RAP2.12 to the promoter containing 5xHRPE for NLUC/FLUC in GG, GP, UG, and UP models | 0.07 | Fitted |
| $HRPE_{1GP}$ | | 0.06 | Fitted |
| $HRPE_{1UG}$ | | 0.04 | Fitted |
| $HRPE_{1UP}$ | | 0.02 | Fitted |
| $HRPE_{2GP}$ | Binding rate of RAP2.12 to the promoter containing 5xHRPE for PCO1 synthesis in GP and UP models | 0.01 | Fitted |
| $HRPE_{2UP}$ | | 0.005 | Fitted |
| $HRPE_{3UG}$ | The binding rate of RAP2.12 to the promoter containing 5xHRPE for RAP2.12 synthesis in UG and UP models | 0.02 | Fitted |
| $HRPE_{3UP}$ | | 0.015 | Fitted |
| $Km_3$ | Michaelis-Menten constant for HRPE activation by RAP2.12 | 0.5 | Fitted |
| $k_5$ | NLUC/FLUC decay rate | 0.01-0.014 | Fitted |
| $k_6$ | PCO1 protein synthesis rate constant measured by RAP2.12 activating $HRPE_2$. | 0.0002-0.003 | Fitted |
| $k_7$ | PCO1 degradation rate | 0.0009 | Fitted |
| $k_8$ | Catalytic rate constant for PCO1-mediated oxidation of RAP2.12 | 9.6688 | Re-fitted (White et al., 2018) |
| $K_{m4}$ | Michaelis-Menten constant for $O_2$ as a substrate of PCO1 | 19.5138 | Re-fitted (White et al., 2018) |
| $K_{m5}$ | Michaelis-Menten constant for RAP2.12 as a substrate of PCO1 | 286.6942 | Re-fitted (White et al., 2018) |
| $k_9$ | RAP2.12 protein synthesis rate constant measured by RAP2.12 activating $HRPE_3$. | 0.03-0.05 | Fitted |

**Table S5. *S. cerevisiae* strains generated in this work.** Name, genotype detailing the expression plasmid's integration site and selection media (Y: YPD; SD-4: SD -his-trp-ura-leu). In bold, wild-type strains. In MaV203, bold indicates the integrated UAS$_{GAL}$ reporters.

| Strain name | Genotype | Media |
|---|---|---|
| **MaV203** | *MATα; leu2-3,112; trp1-901; his3Δ200; ade2-101; $cyh2^R$; $can1^R$; gal4Δ; gal80Δ;* ***GAL1-lacZ; HIS3$_{UASGAL1}$-HIS3@LYS2; SPAL10$_{UASGAL1}$-URA3*** | Y |
| **W3031A** | *MATα; his3-11_15; leu2-3_112; ura3-1; trp1Δ2; ade2-1; can1-100* | Y |
| **W3031B** | *MATα; his3-11_15; leu2-3_112; ura3-1; trp1Δ2; ade2-1; can1-100* | Y |
| MGUS | *MaV203 leu2::LEU2 pAG305GPD-GUS* | SD - leu |
| MPCO | *MaV203 leu2::LEU2 pAG305GPD-PCO4-SV40* | SD - leu |
| MGUS\|R | *MGUS trp1::TRP1 pAG304GPD-RAP2.12* | SD - trp-leu |
| MGUS\|G4S | *MGUS trp1::TRP1 pAG304GPD-GAL4Ste12* | SD - trp-leu |
| MGUS\|SYRAP | *MGUS trp1::TRP1 pAG304GPD-SYRAP* | SD - trp-leu |
| MPCO\|SYRAP | *MPCO trp1::TRP1 pAG304GPD-SYRAP* | SD - trp-leu |
| MGUS\|UbSYRAP | *MGUS trp1::TRP1 pAG304GPD-UbSYRAP* | SD - trp-leu |
| MPCO\|UbSYRAP | *MPCO trp1::TRP1 pAG304GPD- UbSYRAP* | SD - trp-leu |
| MDLOR\|PCO.1 | *MaV203 trp1::TRP1 pAG304GPD-C-DLOR + pAG415GPD-PCO4* | SD - trp-leu |
| MDLOR\|PCO.2 | *MaV203 trp1::TRP1 pAG304GPD-C-DLOR + pAG415GPD-PCO4-SV40* | SD - trp-leu |
| WFLUC | *W3031A his3::HIS3 pAG303GPD-FLUC* | SD - his |
| WH | *WFLUC trp1::TRP1 pAG304GPD-HRPE-NLUC* | SD - trp-his |
| WHO | *WFLUC trp1::TRP1 pAG304GPD-HRPE$_{\Omega}$-NLUC* | SD - trp-his |
| WHA | *WFLUC trp1::TRP1 pAG304GPD-HRPE$_{ADH}$-NLUC* | SD - trp-his |
| WH\|R | *WH ura3::pAG306-RAP2.12* | SD - trp-ura-his |
| WH\|G4S | *WH ura3::pAG306-G4Ste* | SD - trp-ura-his |
| WH\|RG4 | *WH ura3::URA3 pAG306GPD-RAP2.12-G4AD* | SD - trp-ura-his |
| WH\|R6T | *WH ura3::URA3 pAG306GPD-RAP2.12-6TVP* | SD - trp-leu-his |
| WH\|SYRAP | *WH ura3::URA3 pAG306GPD-SYRAP* | SD - trp-ura-his |
| WH\|UbSYRAP | *WH ura3::URA3 pAG306GPD-UbSYRAP* | SD - trp-ura-his |
| WHO\|R | *WHO ura3::pAG306-RAP2.12* | SD - trp-ura-his |
| WHO\|G4S | *WHO ura3::pAG306-G4Ste* | SD - trp-ura-his |
| WHO\|RG4 | *WHO ura3::URA3 pAG306GPD- RAP2.12-G4AD* | SD - trp-ura-his |
| WHO\|R6T | *WHO ura3::URA3 pAG306GPD- RAP2.12-6TVP* | SD - trp-ura-his |
| WHO\|SYRAP | *WHO ura3::URA3 pAG306GPD-SYRAP* | SD - trp-ura-his |
| WHO\|UbSYRAP | *WHO ura3::URA3 pAG306GPD-UbSYRAP* | SD - trp-ura-his |
| WHA\|R | *WHA ura3::pAG306-RAP2.12* | SD - trp-ura-his |
| WHA\|G4S | *WHA ura3::pAG306-G4Ste* | SD - trp-ura-his |
| WHA\|RG4 | *WHA ura3::URA3 pAG306GPD- RAP2.12-G4AD* | SD - trp-ura-his |
| WHA\|R6T | *WHA ura3::URA3 pAG306GPD- RAP2.12-6TVP* | SD - trp-ura-his |
| WHA\|SYRAP | *WHA ura3::URA3 pAG306GPD-SYRAP* | SD - trp-ura-his |
| WHA\|UbSYRAP | *WHA ura3::URA3 pAG306GPD-UbSYRAP* | SD - trp-ura-his |
| WH\|SYRAP\|PCO | *WH\|SYRAP leu2::LEU2 pAG305GPD-PCO4-SV40* | SD-4 |
| WH\|SYRAP\|GUS | *WH\|SYRAP leu2::LEU2 pAG305GPD-GUS* | SD-4 |
| WH\|UbSYRAP\|PCO | *WH\|UbSYRAP leu2::LEU2 pAG305GPD-PCO4-SV40* | SD-4 |
| WH\|UbSYRAP\|GUS | *WH\|UbSYRAP leu2::LEU2 pAG305GPD-GUS* | SD-4 |
| WHA\|SYRAP\|PCO | *WHA\|SYRAP leu2::LEU2 pAG305GPD-PCO4-SV40* | SD-4 |
| WHA\|SYRAP\|GUS | *WHA\|SYRAP leu2::LEU2 pAG305GPD-GUS* | SD-4 |
| WHA\|UbSYRAP\|GUS | *WHA\|UbSYRAP leu2::LEU2 pAG305GPD-GUS* | SD-4 |
| WHA\|UbSYRAP\|PCO **(MynOx)** | *WHA\|UbSYRAP leu2::LEU2 pAG305GPD-PCO4-SV40* | SD-4 |

| **W303 *MATα* strains** | **Genotype** | **Media** |
|---|---|---|
| WB\|H$_A$UbSYRAP | *W3031B his3::HIS3 pAG303HRPE$_{ADH}$-UbSYRAP* | SD-4 |
| WB\|H$_A$GUS | *W3031B his3::HIS3 pAG303HRPE$_{ADH}$-GUS* | SD-4 |
| WB\|H$_A$UbSYHRE | *W3031B his3::HIS3 pAG303HRPE$_{ADH}$-UbSYHRE* | |
| WB\|H$_A$UbSYRAP\|H$_A$PCO | *WB\|H$_A$UbSYRAP leu2::LEU2 Pag305HRPE$_{ADH}$-PCO1-GFP* | SD-4 |

| | | |
|---|---|---|
| WB\|$H_A$UbSYRAP\|$H_A$GUS | *WB\|$H_A$UbSYRAP leu2::LEU2 Pag305HRPE$_{ADH}$-GUS* | SD-4 |
| WB\|$H_A$GUS\|$H_A$PCO | *WB\|$H_A$GUS leu2::LEU2 Pag305HRPE$_{ADH}$-PCO1-GFP* | SD-4 |
| WB\|$H_A$GUS\|$H_A$GUS | *WB\|$H_A$GUS leu2::LEU2 Pag305HRPE$_{ADH}$-GUS* | SD-4 |
| WB\| $H_A$UbSYHRE \|$H_A$PCO | *WB\|$H_A$UbSYHRE leu2::LEU2 Pag305HRPE$_{ADH}$-PCO1* | SD-4 |
| WB\| $H_A$UbSYHRE \|$H_A$GUS | *WB\|$H_A$UbSYHRE leu2::LEU2 Pag305HRPE$_{ADH}$-GUS* | SD-4 |
| **Mated strains** | **Genotype** | **Media** |
| MynOx x UP | *MynOx x WB\|$H_A$UbSYRAP\|$H_A$PCO* | SD-4 |
| MynOx x UG | *MynOx x WB\|$H_A$UbSYRAP\|$H_A$GUS* | SD-4 |
| MynOx x GP | *MynOx x WB\|$H_A$GUS\|$H_A$PCO* | SD-4 |
| MynOx x GG | *MynOx x WB\|$H_A$GUS\|$H_A$GUS* | SD-4 |
| MynOx x HRE2P | *MynOx x WB\| $H_A$UbSYHRE \|$H_A$PCO* | SD-4 |
| MynOx x HRE2G | *MynOx x WB\| $H_A$UbSYHRE \|$H_A$GUS* | SD-4 |

**Table S6. Synthetic sequences used in this study.** Color code for NLUC-UBQ-RAP2.12$_{2-50}$-GAL4STE string: NLUC (Orange), UBQ (Black), RAP2.12 N-terminal fragment corresponding to aa 2-50 (Green), Gal4Ste (Purple for Gal4- and Blue for Ste-derived sequences). For UbSYRAP: UBQ (black), full RAP2.12 coding sequence devoid of initial Met codon (green) and GAL4Ste (purple/blue), flanked by attB sites compatible for BP cloning reaction (red). For ccdB-NLUC: SacI restriction site (bold), attL1 (pink), ccdB cassette with chloramphenicol resistance gene (underlined) and ccdb toxin (light blue), attL2 (pink), NLUC (orange), XhoI restriction site (bold).

| String name | Sequence (5'-3') |
|---|---|
| NLUC-UBQ-RAP2.12$_{2-50}$-GAL4STE | **ATGGTTTTCACCCTTGAGGACTTCGTTGGAGATTGGAGACAGACCGCTGGATACAACCTTGATCAGG**<br>**TGTTGGAGCAAGGTGGGGTGTCATCTTTGTTCCAGAACCTCGGAGTTAGCGTGACCCCTATCCAGAG**<br>**AATCGTTCTCTCTGGTGAGAACGGGCTCAAGATCGATATCCACGTGATCATCCCTTACGAGGGACTT**<br>**AGCGGAGATCAGATGGGACAGATCGAGAAGATTTTCAAGGTGGTGTACCCTGTGGACGACCACCACT**<br>**TCAAGGTTATCCTCCATTACGGAACCCTCGTGATCGATGGTGTGACCCCAAACATGATCGACTACTT**<br>**CGGTAGACCGTACGAGGGAATCGCTGTGTTCGATGGAAAGAAGATTACCGTCACTGGGACCCTCTGG**<br>**AACGGGAACAAGATTATCGATGAGAGGCTCATCAACCCGGACGGCTCACTTTTGTTCAGAGTGACTA**<br>**TCAACGGTGTGACCGGTTGGAGACTTTGCGAGAGAATTTTGGCT**ATGCAAATTTTCGTGAAAACACT<br>CACTGGTAAGACCATCACT**CTCGAG**GTTGAGAGCTCTGACACCATCGACAATGTTAAGGCAAAGATC<br>CAGGACAAGGAAGGCATTCCTCCTGACCAGCAAAGATTGATCTTCGCTGGTAAGCAGCTAGAAGATG<br>GCCGCACCTTGGCTGATTACAACATCCAGAAAGAATCAACACTCCACTTGGTTCTCAGGTTAAGAGG<br>TGGTTGTGGGGGAGCTATCATTTCTGATTTCATCTGGTCGAAATCTGAGTCAGAACCGAGTCAACTC<br>GGCTCTGTTAGCAGCAGGAAGAAGCGTAAACCCGTCTCAGTGAGTGAAGAAAGAGATGGGAAACGAG<br>AGAGGAAGAATCTGTAC**ATGAAGCTGTTGTCCTCTATTGAACAAGCTTGCGATATCTGCAGGCTGAA**<br>**AAAATTGAAGTGCTCCAAAGAAAAGCCAAAGTGTGCTAAGTGTTTGAAGAACAACTGGGAATGTAGA**<br>**TACTCTCCAAAGACTAAGAGATCTCCATTGACTAGAGCACATTTGACCGAAGTTGAATCTAGGTTGG**<br>**AAAGATTGGAGCAGCTGTTCTTGTTGATTTTCCCAAGAGAAGATCTGGACATGATCTTGAAGATGGA**<br>**TTCCTTGCAAGATATCAAGGCTTTGTTGACTGGTTTGTTCGTTCAAGATAACGTTAACAAGGATGCC**<br>**GTTACTGATAGATTGGCTTCTGTTGAAACTGATATGCCATTGACCTTGAGACAACATAGAATTTCTG**<br>**CTACCTCCAGCTCTGAAGAATCTTCTAACAAAGGTCAAAGACAGTTGACCGTGTCTAGAGATGAAGA**<br>**AGATTTTCCATTGGATTACTTCCCAGTCTCTGTTGAATATCCAACTGAAGAAAATGCCTTCGATCCA**<br>**TTTCCACCACAAGCTTTTACTCCAGCTATTGATTCTGCTGCTCATCATGATAACTCTACTATCCCAT**<br>**TAGACTTCATGCCAAGAGATGCATTGCATGGTTTTGATTGGTCTGAAGAGGATGATATGTCTGATGG**<br>**TTTGCCATTCTTGAAAACCGATCCAAACAACAACGGTTTCTAA** |
| UbSYRAP | GGGGACAAGTTTGTACAAAAAAGCAGGCTCCATGCAAATTTTCGTGAAAACACTCACTGGTAAGACC<br>ATCACT**CTCGAG**GTTGAGAGCTCTGACACCATCGACAATGTTAAGGCAAAGATCCAGGACAAGGAAG<br>GCATTCCTCCTGACCAGCAAAGATTGATCTTCGCTGGTAAGCAGCTAGAAGATGGCCGCACCTTGGC<br>TGATTACAACATCCAGAAAGAATCAACACTCCACTTGGTTCTCAGGTTAAGAGGTGGTTGTGGAGGA<br>GCTATAATATCCGATTTCATTCCACCGCCGAGGTCTCGCCGTGTTACTAGCGAGTTTATTTGGCCGG<br>ATCTGAAGAAGAATTTGAAAGGATCGAAGAAAAGCTCGAAGAATCGTTCGAATTTCTTCGATTTTGA<br>CGCTGAGTTCGAAGCTGATTTCCAAGGTTTCAAAGATGATTCGTCTATCGATTGCGATGATGATTTC<br>GACGTCGGTGATGTTTTCGCCGATGTGAAACCATTCGTTTTCACTTCGACTCCAAAACCCGCCGTCT<br>CCGCCGCTGCGGAAGGTTCAGTTTTTGGTAAGAAAGTTACTGGCTTGGATGGGGACGCTGAGAAATC<br>TGCAAATAGGAAGAGGAAGAATCAGTACCGAGGGATTAGGCAACGTCCTTGGGGAAAATGGGCTGCT<br>GAGATACGTGATCCAAGGGAAGGTGCTAGAATCTGGCTTGGAACGTTCAAGACAGCTGAGGAAGCTG<br>CTAGAGCTTACGATGCTGCAGCGCGGAGAATCCGTGGATCTAAAGCTAAGGTGAATTTCCCTGAAGA<br>AAACATGAAGGCTAATTCTCAGAAACGCTCTGTGAAGGCTAATCTTCAGAAACCAGTGGCTAAACCT<br>AACCCTAACCCAAGTCCAGCTTTGGTTCAGAACTCGAACATCTCCTTTGAAAATATGTGTTTCATGG<br>AGGAGAAACACCAAGTGAGCAACAACAACAACAACCAGTTTGGGATGACAAACTCCGTTGATGCTGG<br>ATGTAATGGGTATCAGTATTTCAGCTCTGACCAGGGTAGTAATTCTTTCGATTGTTCGGAGTTTGGT<br>TGGAGCGATCAAGCTCCGATAACTCCCGACATCTCTTCTGCGGTTATCAACAACAACAACTCAGCTC<br>TGTTCTTTGAGGAAGCCAATCCAGCTAAGAAGCTCAAGTCTATGGATTTCGAGACACCTTACAACAA<br>CACTGAATGGGACGCTTCACTGGATTTCCTCAACGAAGATGCTGTAACGACTCAGGACAATGGTGCA<br>AACCCTATGGACCTATGGAGTATTGATGAAATTCATTCCATGATTGGAGGAGTCTTC**ATGAAGCTGT**<br>**TGTCCTCTATTGAACAAGCTTGCGATATCTGCAGGCTGAAAAAATTGAAGTGCTCCAAAGAAAAGCC**<br>**AAAGTGTGCTAAGTGTTTGAAGAACAACTGGGAATGTAGATACTCTCCAAAGACTAAGAGATCTCCA**<br>**TTGACTAGAGCACATTTGACCGAAGTTGAATCTAGGTTGGAAAGATTGGAGCAGCTGTTCTTGTTGA**<br>**TTTTCCCAAGAGAAGATCTGGACATGATCTTGAAGATGGATTCCTTGCAAGATATCAAGGCTTTGTT**<br>**GACTGGTTTGTTCGTTCAAGATAACGTTAACAAGGATGCCGTTACTGATAGATTGGCTTCTGTTGAA**<br>**ACTGATATGCCATTGACCTTGAGACAACATAGAATTTCTGCTACCTCCAGCTCTGAAGAATCTTCTA**<br>**ACAAAGGTCAAAGACAGTTGACCGTGTCTAGAGATGAAGAAGATTTTCCATTGGATTACTTCCCAGT**<br>**CTCTGTTGAATATCCAACTGAAGAAAATGCCTTCGATCCATTTCCACCACAAGCTTTTACTCCAGCT**<br>**ATTGATTCTGCTGCTCATCATGATAACTCTACTATCCCATTAGACTTCATGCCAAGAGATGCATTGC**<br>**ATGGTTTTGATTGGTCTGAAGAGGATGATATGTCTGATGGTTTGCCATTCTTGAAAACCGATCCAAA**<br>**CAACAACGGTTTCTAA**cacccagctttcTTGTACAAAGTGGTCCCC |

| | |
|---|---|
| ccdB-NLUC | **gagctcACAAGTTTGTACAAAAAAGCTGAACGAGAAACGTAAAATGATATAAATATCAATATATTAA**<br>**ATTAGATTTTGCATAAAAAACAGACTACATAATACTGTAAAACACAACATATCCAGTCACTATG**GCG<br>GCCGCATTAGGCACCCCAGGCTTTACACTTTATGCTTCCGGCTCGTATAATGTGTGGATTTTGAGTT<br>AGGATCCGGCGAGATTTTCAGGAGCTAAGGAAGCTAAAATGGAGAAAAAAATCACTGGATATACCAC<br>CGTTGATATATCCCAATGGCATCGTAAAGAACATTTTGAGGCATTTCAGTCAGTTGCTCAATGTACC<br>TATAACCAGACCGTTCAGCTGGATATTACGGCCTTTTTAAAGACCGTAAAGAAAAATAAGCACAAGT<br>TTTATCCGGCCTTTATTCACATTCTTGCCCGCCTGATGAATGCTCATCCGGAATTCCGTATGGCAAT<br>GAAAGACGGTGAGCTGGTGATATGGGATAGTGTTCACCCTTGTTACACCGTTTTCCATGAGCAAACT<br>GAAACGTTTTCATCGCTCTGGAGTGAATACCACGACGATTTCCGGCAGTTTCTACACATATATTCGC<br>AAGATGTGGCGTGTTACGGTGAAAACCTGGCCTATTTCCCTAAAGGGTTTATTGAGAATATGTTTTT<br>CGTCTCAGCCAATCCCTGGGTGAGTTTCACCAGTTTTGATTTAAACGTGGCCAATATGGACAACTTC<br>TTCGCCCCCGTTTTCACCATGGGCAAATATTATACGCAAGGCGACAAGGTGCTGATGCCGCTGGCGA<br>TTCAGGTTCATCATGCCGTCTGTGATGGCTTCCATGTCGGCAGAATGCTTAATGAATTACAACAGTA<br>CTGCGATGAGTGGCAGGGCGGGGCGTAAACGCGTGGATCCGGCTTACTAAAAGCCAGATAACAGTAT<br>GCGTATTTGCGCGCTGATTTTTGCGGTATAAGAATATATACTGATATGTATACCCGAAGTATGTCAA<br>AAAGAGGTGTGCTATGAAGCAGCGTATTACAGTGACAGTTGACAGCGACAGCTATCAGTTGCTCAAG<br>GCATATATGATGTCAATATCTCCGGTCTGGTAAGCACAACCATGCAGAATGAAGCCCGTCGTCTGCG<br>TGCCGAACGCTGGAAAGCGGAAAATCAGGAAGGGATGGCTGAGGTCGCCCGGTTTATTGAAATGAAC<br>GGCTCTTTTGCTGACGAGAACAGGGACTGGTGAAATGCAGTTTAAGGTTTACACCTATAAAAGAGAG<br>AGCCGTTATCGTCTGTTTGTGGATGTACAGAGTGATATTATTGACACGCCCGGGCGACGGATGGTGA<br>TCCCCCTGGCCAGTGCACGTCTGCTGTCAGATAAAGTCTCCCGTGAACTTTACCCGGTGGTGCATAT<br>CGGGGATGAAAGCTGGCGCATGATGACCACCGATATGGCCAGTGTGCCGGTCTCCGTTATCGGGGAA<br>GAAGTGGCTGATCTCAGCCACCGCGAAAATGACATCAAAAACGCCATTAACCTGATGTTCTGGGGAA<br>TATAAATGTCAGGCTCCCTTATACACAGCCAGTCTGCAGGTCGAC**CATAGTGACTGGATATGTTGTG**<br>**TTTTACAGTATTATGTAGTCTGTTTTTTATGCAAAATCTAATTTAATATATTGATATTTATATCATT**<br>**TTACGTTTCTCGTTCAGCTTTCTTGTACAAAGTGG**TGATAAAAAA**ATGGTTTTCACCCTTGAGGACT**<br>**TCGTTGGAGATTGGAGACAGACCGCTGGATACAACCTTGATCAGGTGTTGGAGCAAGGTGGGGTGTC**<br>**ATCTTTGTTCCAGAACCTCGGAGTTAGCGTGACCCCTATCCAGAGAATCGTTCTCTCTGGTGAGAAC**<br>**GGGCTCAAGATCGATATCCACGTGATCATCCCTTACGAGGGACTTAGCGGAGATCAGATGGGACAGA**<br>**TCGAGAAGATTTTCAAGGTGGTGTACCCTGTGGACGACCACCACTTCAAGGTTATCCTCCATTACGG**<br>**AACCCTCGTGATCGATGGTGTGACCCCAAACATGATCGACTACTTCGGTAGACCGTACGAGGGAATC**<br>**GCTGTGTTCGATGGAAAGAAGATTACCGTCACTGGGACCCTCTGGAACGGGAACAAGATTATCGATG**<br>**AGAGGCTCATCAACCCGGACGGCTCACTTTTGTTCAGAGTGACTATCAACGGTGTGACCGGTTGGAG**<br>**ACTTTGCGAGAGAATTTTGGCTTAActcgag** |

**Table S7. Cloning primers used in this work.** In bold, adaptors for BP cloning.

| Function | Primer name | Sequence (5'-3') |
|---|---|---|
| PCO4-SV40 cloning | PCO4_BP_Fw | **AAAAAAGCAGGCTCC**ATGCCTTACTTTGCT |
| | PCO4_SV40_Rv1 | CACTTTGCGTTTTTTTTTCGGAGTTCTAATGACA |
| | PCO4SV40_BP_Rv2 | **AGAAAGCTGGGTG**TTACACTTTGCGTTTTT |
| RAP2.12 | R2.12_Fw | **AAAAAAGCAGGCTCC**ATGTGTGGAGGAGCTA |
| RAP2.12-G4AD overlapping | OL_RAP_G4AD_Fw | ATTGGAGGAGTCTTCGCCAATTTTAATCAA |
| | OL_RAP_G4AD_Rv | TTGATTAAAATTGGCGAAGACTCCTCCAAT |
| | G4AD_BP_Rv | **AGAAAGCTGGGTG**TCACTCTTTTTTTGGGTTTG |
| RAP2.12-6TVP overlapping | OL_RAP_6TVP_Fw | GATTGGAGGAGTCTTCGGTGGTGGATCTGGTG |
| | OL_RAP_6TVP_Rv | CACCAGATCCACCACCGAAGACTCCTCCAATC |
| | 6TVPSV40_BP_Rv | **AGAAAGCTGGGTG**TCATTACACTTTGCGTTTTTTTTTCGG |
| SYRAP overlapping | OL_R2.12-G4Ste_Fw | ATTGGAGGAGTCTTCATGAAGCTGTTGTCC |
| | OL_G4Ste – R2.12_Rv | GGACAACAGCTTCATGAAGACTCCTCCAAT |
| | G4Ste_Rv | **AGAAAGCTGGGTG**TTAGAAACCGTTGTTGTTT |
| UbSYRAP overlapping | SY_UBQ_Fw | **AAAAAAGCAGGCTCC**ATGCAAATTTTCGTGAAA |
| | OL_SY_Fw | AGGTTAAGAGGTGGTTGTGGAGGAGCTATA |
| | OL_SY_Rv | TATAGCTCCTCCACAACCACCTCTTAACCT |
| PCO1-GFP overlapping | PCO1gwFw | **AAAAAAGCAGGCTCC**ATGGGGTTTGAGATGAAACC |
| | PCO1GFP_OL_Fw | CCAAAGGTTGAAGATatggtgagcaagggc |
| | PCO1GFP_OL_Rv | gcccttgctcaccatATCTTCAACCTTTGG |
| | PCO1gwRv | **AGAAAGCTGGGTG**tcacttgtacagctcgtccatgc |
| UbSYHRE overlapping | UbSYHRE_UBgwFw | **AAAAAAGCAGGCTCC**ATGCCAACTTTGTACAAAAA |
| | UbSYHRE_HRG4OL_Fw | CAAGACTCCAATTAAATGAAGCTGTTGTCC |
| | UbSYHRE_HRG4OL_Rv | GGACAACAGCTTCATTTAATTGGAGTCTTG |
| | UbSYHRE_G4SgwRv | **AGAAAGCTGGGTG**TTAGAAACCGTTGTTGTTTGGAT |
| pAG303- and pAG305-HRPE$_{ADH}$ cloning | HRPE5xA30x_HF | GGCGAATTGGAGCTCTTCGAGTTTACCGCGGCCGCCCCCT |
| | HRPE5xA30x _p30xR | ATTCAGGTGAAGGGGGCGGCCGCGGTAAACTCGAAGAGCT |
| | HRPE30x_30xRv | CGTCGATCAATGTTGGATATCTAAACTCGAAGA |
| | HRPE5xA30x_p30xF | AAAAGCAAGTTCTTCACTGTTGATACGGATTCTAGAACTA |
| AttB1 adaptors | BP_Ad_Fw | GGGGACAAGTTTGTACAAAAAAGCAGGCT |
| | BP_Ad_Rv | GGGGACCACTTTGTACAAGAAAGCTGGGT |

**Table S8. Gateway-compatible vectors from this work.** Inserts were recombined into the appropriate destination vectors from the list, as indicated in the main text, to generate the desired expression vectors. Descriptions of the destination vectors include auxotrophic markers, in the case of yeast expression plasmids, or selective markers for Agrobacteria/plants in the case of plant expression plasmids, and the features of the transgene expression cassette.

| **Entry vector** | **Insert** | **Source** |
|---|---|---|
| HRPE_pE | 5xHRPE | (13) |
| $HRPE_{\Omega}$_pE | 5xHRPE-35SΩ leader | (13) |
| $HRPE_{ADH}$_pE | 5xHRPE-ADH 5' UTR | (13) |
| GUS_pE | β-glucoronidase (GUS) | Thermo-Fisher Sci. |
| RAP2.12_pE | RAP2.12 | (14) |
| PCO4_pE | PCO4 | (2) |
| C-DLOR_pE | RLUC-Ub-$RAP2.12_{2\text{-}28}$-FLUC | (2) |
| PCO4-SV40_201 | PCO4 fused to the SV40 NLS | This study |
| G4Ste_201 | $GAL4_{1\text{-}147}$-$STE12_{301\text{-}335}$Y310F-$GAL4_{148\text{-}196}$ | This study |
| IB+_201 | NLUC-AtUBQ4-$RAP2.12_{1\text{-}50}$-Y310F$GAL4_{1\text{-}147}$-$Ste12_{301\text{-}335}$-$GAL4_{148\text{-}196}$ | This study |
| SYRAP_201 | RAP2.12-$GAL4_{1\text{-}147}$-$Ste12_{301\text{-}335}$Y310F-$GAL4_{148\text{-}196}$ | This study |
| RAP2.12-G4AD_201 | RAP2.12-GAL4AD | This study |
| RAP2.12-6TVP_201 | RAP2.12-6XTAL-4XVP16 | This study |
| UbSYRAP_201 | AtUBQ4-$RAP2.12_{2\text{-}358}$-$GAL4_{1\text{-}147}$-$Ste12_{301\text{-}335}$Y310F-$GAL4_{148\text{-}196}$ | This study |
| UbSYHRE_201 | AtUBQ4-$HRE2_{2\text{-}171}$-$GAL4_{1\text{-}147}$-$Ste12_{301\text{-}335}$Y310F-$GAL4_{148\text{-}196}$ | This study |
| **Destination vector** | **Description** | **Source** |
| pAG304GPD | Integrative plasmid TRP1, pGPD-ccdb-TCYC1 | Addgene (#14135) |
| pAG305GPD | Integrative plasmid LEU2, pGPD-ccdb-TCYC1 | Addgene (#14138) |
| pAG306GPD | Integrative plasmid URA3, pGPD-ccdb-TCYC1 | Addgene (#14140) |
| pAG303GPD | Integrative plasmid HIS3, pGPD-ccdb-TCYC1 | Addgene (#14134) |
| pAG304NLUC | Integrative plasmid TRP1, ccdb-NLUC-TCYC1 | This study |
| pK7WG2 | Plant binary vector (Strep-Spec/Kan), p35S-ccdb-t35S | (4) |
| pBGWL7 | Plant binary vector (Strep-Spec/Kan), ccdb-t35S | (4) |
| pAG303_$HRPE_{ADH}$ | Integrative plasmid HIS3, HRPEADH-ccdb-TCYC1 | This study |
| pAG305_$HRPE_{ADH}$ | Integrative plasmid LEU2, HRPEADH-ccdb-TCYC1 | This study |
| pBGWNLUCL7 | Plant binary vector (Strep-Spec/Kan), ccdb-NLUC-t35S | This study |

**Table S9. qPCR primers used in this work.** For the wild-type genes analyzed, the Arabidopsis Genome Initiative (AGI) identifier, or the yeast systematic code are given as codes.

| Gene and code | Primer name | Sequence (5'-3') |
|---|---|---|
| AtUBQ10 (AT4G05320) | AtUBQ10_F | GGCCTTGTATAATCCCTGATGAATAAG |
| | AtUBQ10_R | AAAGAGATAACAGGAACGGAAACATAGT |
| AtADH1 (AT1G77120) | AtADH1_F | TATTCGATGCAAAGCTGCTGTG |
| | AtADH1_R | CGAACTTCGTGTTTCTGCGGT |
| AtPDC1 (AT4G33070) | AtPDC1_F | CACAGAATCTTCAATGCTTCTTACC |
| | AtPDC1_R | CCATGATAAAGCGTACATGGAA |
| AtPGB1 (AT2G16060) | AtPGB1_F | TTTGAGGTGGCCAAGTATGCA |
| | AtPGB1_R | TGATCATAAGCCTGACCCCAA |
| AtHRE2 (AT2G47520) | AtHRE2_F | GAAGCGTAAACCCGTCTCAGTG |
| | AtHRE2_R | TTTGCTCGGGYCACGAATCT |
| AtLBD41 (AT3G02550) | AtLBD41_F | TGAAGCGCAAGCTAACGCA |
| | AtLBD41_R | ATCCCAGGACGAAGGTGATTG |
| AtPCO1 (AT5G15120) | AtPCO1_F | ATTGGGTGGTTGATGCTCCAATG |
| | AtPCO1_R | ATGCATGTTCCCGCCATCTTC |
| AtSAD6 (AT1G43800) | AtSAD6_F | TTGTGGAAGGTGAAGGCAAGGG |
| | AtSAD6_R | TTGGCAACCCGCTTCTTTCTTACC |
| AtSUS4 (AT3G43190) | AtSUS4_F | CGCAGAACGTGTAATAACGCG |
| | AtSUS4_R | CAACCCTTGAGAGCAAAGCAAA |
| AtHRA1 (AT3G10040) | AtHRA1_F | ACAACCACCGCAACAGAATCC |
| | AtHRA1_R | TCTCCGCAATTCTCGCCAT |
| NLUC | NLUC_F | CCAGAACCTCGGAGTTAGCGTG |
| | NLUC_R | TCTGTCCCATCTGATCTCCGCT |
| ScACT1 (YFL039C) | ScACT1_F | CCATCCAAGCCGTTTTGTCC |
| | ScACT1_R | GGCGTGAGGTAGAGAGAAACC |
| UbSYRAP | UbSYRAP_F | AATGGGACGCTTCACTGGATTTC |
| | UbSYRAP_R | ATCGCAAGCTTGTTCAATAGAG |
| ScROX1 (YPR065W) | ScROX1_F | GTCCACAACTACCCCTACGC |
| | ScROX1_R | TAGCGGTGACCTCAGTGTTG |
| ScHEM13 (YDR044W) | ScHEM13_F | CACCAACTGCACAAGGATGC |
| | ScHEM13_R | CGTGTTTCCTTACGGTGGGT |
| ScCOX5B (YIL111W) | ScCOX5B_F | GATCTGCCCGAAAGATGGGAA |
| | ScCOX5B_R | CCACTCGCCGTAGGATATGT |
| ScFRT2 (YAL028W) | ScFRT2_F | TCGCATTCATCAGACGGGAG |
| | ScFRT2_R | GGCAGGAGAGGATTCGGAAG |

**Table S10. Statistical analysis supporting Fig. S6D.** Student's t tests were performed on UbSYRAP/TUB band intensity and discovery determined by the Two-stage linear step-up procedure of Benjamini, Krieger and Yekutieli. P values of the analysis are displayed (in bold, p values < 0.05).

| | UP-UG | UP-GP | UP-GG | UG-GP | UG-GG | GP-GG |
|---|---|---|---|---|---|---|
| 0' | 0,528 | 0,557 | 0,336 | 0,717 | 0,969 | 0,526 |
| 10' | 0,117 | 0,162 | 0,105 | **0,062** | 0,055 | 0,484 |
| 30' | **0,017** | **0,011** | 0,051 | **0,005** | **0,022** | 0,911 |
| 60' | **0,066** | **0,052** | **0,015** | **0,003** | **0,001** | **0,037** |

**Table S11. Statistical analysis supporting Fig. S8.** For each gene, Student's t tests were performed and discovery determined by the Two-stage linear step-up procedure of Benjamini, Krieger and Yekutieli. P values of the analysis are displayed (in bold, p<0.05).

| *NLUC* | UP-UG | UP-GP | UP-GG | UG-GP | UG-GG | GP-GG |
|---|---|---|---|---|---|---|
| 0' | **0,003** | 0,093 | **0,013** | **0,002** | 0,075 | **0,004** |
| 5' | **0,009** | **0,001** | 0,483 | **0,000** | 0,105 | **0,018** |
| 10' | 0,388 | **0,009** | 0,199 | **0,001** | 0,579 | **0,000** |
| 15' | 0,938 | **0,008** | 0,217 | **0,000** | 0,076 | **0,001** |
| 30' | 0,548 | **0,000** | **0,002** | **0,005** | **0,037** | **0,000** |
| 60' | 0,498 | **0,000** | **0,005** | **0,000** | **0,005** | 0,083 |
| 2h | 0,796 | **0,003** | **0,013** | **0,001** | **0,006** | **0,000** |
| 4h | 0,639 | **0,001** | **0,006** | **0,000** | **0,001** | **0,000** |
| *PCO1* | UP-UG | UP-GP | UP-GG | UG-GP | UG-GG | GP-GG |
| 0' | **0,039** | 0,198 | | 0,055 | | |
| 5' | **0,000** | **0,000** | | **0,000** | | |
| 10' | **0,021** | **0,029** | | **0,034** | | |
| 15' | **0,018** | **0,024** | | **0,023** | | |
| 30' | **0,000** | **0,000** | | **0,010** | | |
| 60' | **0,001** | **0,001** | | **0,001** | | |
| 2h | **0,001** | **0,002** | | **0,019** | | |
| 4h | **0,035** | **0,041** | | **0,003** | | |
| *UbSYRAP* | UP-UG | UP-GP | UP-GG | UG-GP | UG-GG | GP-GG |
| 0' | **0,003** | 0,093 | **0,013** | **0,002** | 0,075 | **0,004** |
| 5' | **0,009** | **0,001** | 0,483 | **0,000** | 0,105 | **0,018** |
| 10' | 0,388 | **0,009** | 0,199 | **0,001** | 0,579 | **0,000** |
| 15' | 0,938 | **0,008** | 0,217 | **0,000** | 0,076 | **0,001** |
| 30' | 0,548 | **0,000** | **0,002** | **0,005** | **0,037** | **0,000** |
| 60' | 0,498 | **0,000** | **0,005** | **0,000** | **0,005** | 0,083 |
| 2h | 0,796 | **0,003** | **0,013** | **0,001** | **0,006** | **0,000** |
| 4h | 0,639 | **0,001** | **0,006** | **0,000** | **0,001** | **0,000** |
